# The Median Preoptic Nucleus is a Key Site for Estradiol Regulation of Sleep-Wake Behaviors in Females Rats

**DOI:** 10.1101/2025.11.06.687021

**Authors:** Katie Kruk, Philip C. Smith, Danielle M. Cusmano, Shaun S. Viechweg, Carissa Byrd, Amy Huddleson, Michael D. Schwartz, Jessica A. Mong

## Abstract

**Study Objectives:** Women experience more sleep disruptions than men, particularly during hormonal transitions such as puberty, pregnancy, and menopause. This study investigated the role of estradiol (E2) in regulating sleep-wake behavior in female rats and identified the brain regions involved.

**Methods:** Using an exogenous E2 replacement model in ovariectomized rats, we assessed changes in sleep-wake patterns via EEG/EMG telemetry. The effects of E2 and progesterone, selective estrogen receptor agonists, and direct brain infusions of E2 and receptor antagonists were evaluated.

**Results:** E2 administration increased wakefulness, reduced non-rapid eye movement (NREM) and rapid eye movement (REM) sleep, and decreased NREM slow-wave activity (SWA), predominantly during the dark phase. These effects required both estrogen receptor alpha (ERα) and beta (ERβ) activation and were mediated by estrogen receptor signaling within the median preoptic nucleus (MnPO). Direct infusion of E2 into the MnPO was sufficient to replicate systemic effects, while local infusion of the pure estrogen receptor antagonist ICI 182,780 (Fulvestrant) attenuated them. Progesterone did not augment E2’s actions, and males showed no sleep-wake changes in response to E2, highlighting sex-specific mechanisms.

**Conclusions:** The MnPO is a critical site where E2 regulates sleep-wake behavior. These findings provide a neurobiological framework for understanding how ovarian hormones contribute to sleep disruptions in women, offering potential therapeutic targets for sleep disorders related to hormonal changes.

**Statement of significance:** Sleep disturbances disproportionately affect women, particularly during hormonal transitions such as puberty, pregnancy, and menopause. Despite this, the underlying mechanisms by which estradiol (E2) influences sleep-wake behavior have remained unclear. This study identifies the median preoptic nucleus (MnPO) as a critical site where E2 signaling regulates sleep-wake patterns in female rats. Our findings demonstrate that E2 increases wakefulness and reduces both non-rapid eye movement (NREM) and rapid eye movement (REM) sleep, with effects primarily observed during the dark phase. By showing that local E2 signaling in the MnPO is both necessary and sufficient for these changes, this work offers a novel neurobiological framework for understanding how fluctuations in ovarian hormones contribute to sleep disruptions in women. These insights lay the groundwork for developing targeted therapies to improve sleep health in women, particularly those experiencing hormonal changes.

## Introduction

Quality sleep is imperative for the maintenance of good health. There is a growing recognition of sex disparities in sleep and rhythm disorders. Compared to males, females are twice as likely to experience sleep disruptions and insomnia throughout their lifespan.^1^ Chronic insufficient sleep is a risk factor for a variety of psychological^2–6^, neurological^7–12^, and neurodegenerative pathologies^9^ as well as cardiovascular and metabolic dysfunctions.^13–17^ Clinical studies reveal that women suffering from sleep disturbances and insufficient sleep are at greater risk compared to men for mood disorders such as depression^18^ as well as metabolic^19^ and cardiovascular dysfunction.^16,20–22^ Yet, historically, the female physiology has been underrepresented in preclinical and clinical studies of sleep and its disorders.

While ovarian steroids, biological sex, and gender are implicated as risk factors for sleep disruptions and insomnia, the exact influence of ovarian steroids on the sleep-wake neurocircuitry remains a significant gap in our knowledge. The increased risks for sleep disturbances in women emerge at the time of puberty and have been linked to fluctuations in the gonadal steroid milieu.^23^ Recent data from women of reproductive age suggest that the presence of estrogens are a contributing factor to sleep disturbances. By contrast, in women approaching menopause, the loss of estrogens is associated with insomnia, frequent night-time awakenings and poor sleep.^24–28^ Thus, a better understanding of how estrogens are influencing the sleep-circuitry may help explain the biological basis for these apparent paradoxical effects of estrogens on sleep disturbances across a women’s lifespan.

Ovarian steroid effects on sleep have been investigated in numerous rodent studies over the past 60 years. While variations in rodent species and strains, steroid dosages, and sleep-wake quantification methods have led to some mixed conclusions about the exact role of ovarian steroids in sleep regulation, the major findings support that estradiol (E2) reduces time spent in sleep states (non-rapid eye movement, NREM and/or REM) with a concomitant increase in the time spent in the wakefulness during the rodent’s active phase.^29–42^ Building on this body work, our previous work, demonstrates (1) the absence of circulating gonadal steroids eliminates sex differences in sleep behavior and architecture and (2) males do not exhibit such fluctuations in response to changes in circulating testosterone or E2.^40^ Taken together, these findings suggest that sex differences in sleep are primarily dependent on the steroidal modulation of sexually differentiated sleep circuitry.^1,40^ Nevertheless, to date, few studies have explored the underlying mechanisms by which ovarian steroids influence sleep.

We have identified sex differences in neuronal activity within the ventrolateral preoptic area (VLPO), a well-established sleep-regulating nucleus.^33,40^ Our recent research also indicates that the median preoptic area (MnPO), another key sleep-regulating nucleus in rats (for review^43,44^), is sensitive to the effects of E2, with E2 influencing adenosine signaling in the MnPO of adult female rats.^42^ In the current study, we sought to expand our knowledge of the mechanisms through which ovarian steroids regulate sleep-wake behavior in adult female rats.

We have previously validated an exogenous E2 replacement model in ovariectomized rats that mimics the proestrus E2 surge of the rat estrous cycle and reproduces the sleep-wake patterns typically observed in females with cycling ovarian hormones.^28,33,38,40,45^ Using this model, we tested whether progesterone, which also surges sharply in the afternoon of proestrus following the E2 peak, enhances or counteracts E2’s effect on sleep-wake behavior and sleep homeostasis. Our findings confirmed that E2 alone was sufficient to change sleep-wake patterns. Next, we addressed whether the E2 effects are driven by classical estrogen receptor actions via estrogen receptor alpha (ERα; product of the *ESR1* gene) and estrogen receptor beta (ERβ; product of the *ESR2* gene). We found that both receptor subtypes are required for the full effect on sleep-wake behavior. Additionally, we report here that ERα protein and mRNA expression of the *ESR1* and *ESR2* genes are localized to the MnPO. Finally, these findings led us to investigate whether local estrogenic signaling in the MnPO is necessary and sufficient to induce changes in sleep-wake behavior. Direct infusion of E2 into the MnPO was sufficient to recapitulate the sleep-wake changes of global E2 treatment, while local infusion of a pure estrogen receptor antagonist, ICI 182,780 (Fulvestrant) into the MnPO blocked such effects. The current findings strongly suggest that that E2 action in the MnPO is necessary and sufficient to induce changes in sleep-wake behavior in the female rat.

## Material and Methods

### Animals

Adult female and male Sprague–Dawley (250-400g) were purchased from Charles River Laboratories (Kingston, N.Y.) and housed in the laboratory animal facilities at the University of Maryland, School of Medicine under a reversed 12 h:12 h dark:light cycle with food and water available ad libitum. In all experiments, zeitgeber time 0 (ZT 0) represents lights ON except where noted. All experimental procedures were run in multiple cohorts, with all experimental groups represented in each cohort. All procedures were performed in accordance with the National Institutes of Health guide for care and use of laboratory animals. All experiments were approved by and were in accordance with the guidelines of the University of Maryland Institutional Animal Care and Use Committee.

### General Surgical Procedures

#### Gonadectomy

All surgeries were conducted under isoflurane anesthesia. An oxygen/isoflurane mixture was delivered through a precision vaporizer. Animals were place in a chamber and loss of consciousness was induced at 3-5% isoflurane. During surgery, animals were maintained with 1-3% delivered through a nose cone. All female rats (250-300g) were ovariectomized (OVX). If males (350-400g) were used for sex difference comparisons, they were either orchiectomized (ORDX) or left gonadally intact in a sham ORDX. Briefly, OVX was performed using a dorsal incision followed by isolation, ligation and removal of the ovaries bilaterally. For male ORDX, an incision was made into the scrotum followed by a small incision into the muscle. The blood supply to each testis was ligated and both testes were removed. The muscle wall and skin were both sutured closed.

#### EEG/EMG Telemetry

At the time of gonadectomy, animal designated for sleep-wake recordings were implanted with TL11M2-F40-EET transmitters (Data Sciences International, St. Paul, MN) according to our standard protocols. Following the placement of the transmitter, another incision was made along the midline of the head and neck to expose the skull and neck muscle. Two burr holes were drilled asymmetrically and stainless-steel screws (Plastics One, Roanoke, VA.) were implanted at 2 mm anterior/1.5 mm lateral and 7 mm posterior/1.5 mm lateral relative to the bregma. The four transmitter leads were threaded subcutaneously to the head incision. Electroencephalographic (EEG) leads were wrapped around the screws and secured with dental cement. Electromyogram (EMG) leads were implanted directly in the dorsal cervical neck muscle, approximately 1.0 mm apart, and sutured in place. Figure 1 presents representative EEG/EMG recording from the described electrode placements.

**Figure 1.**
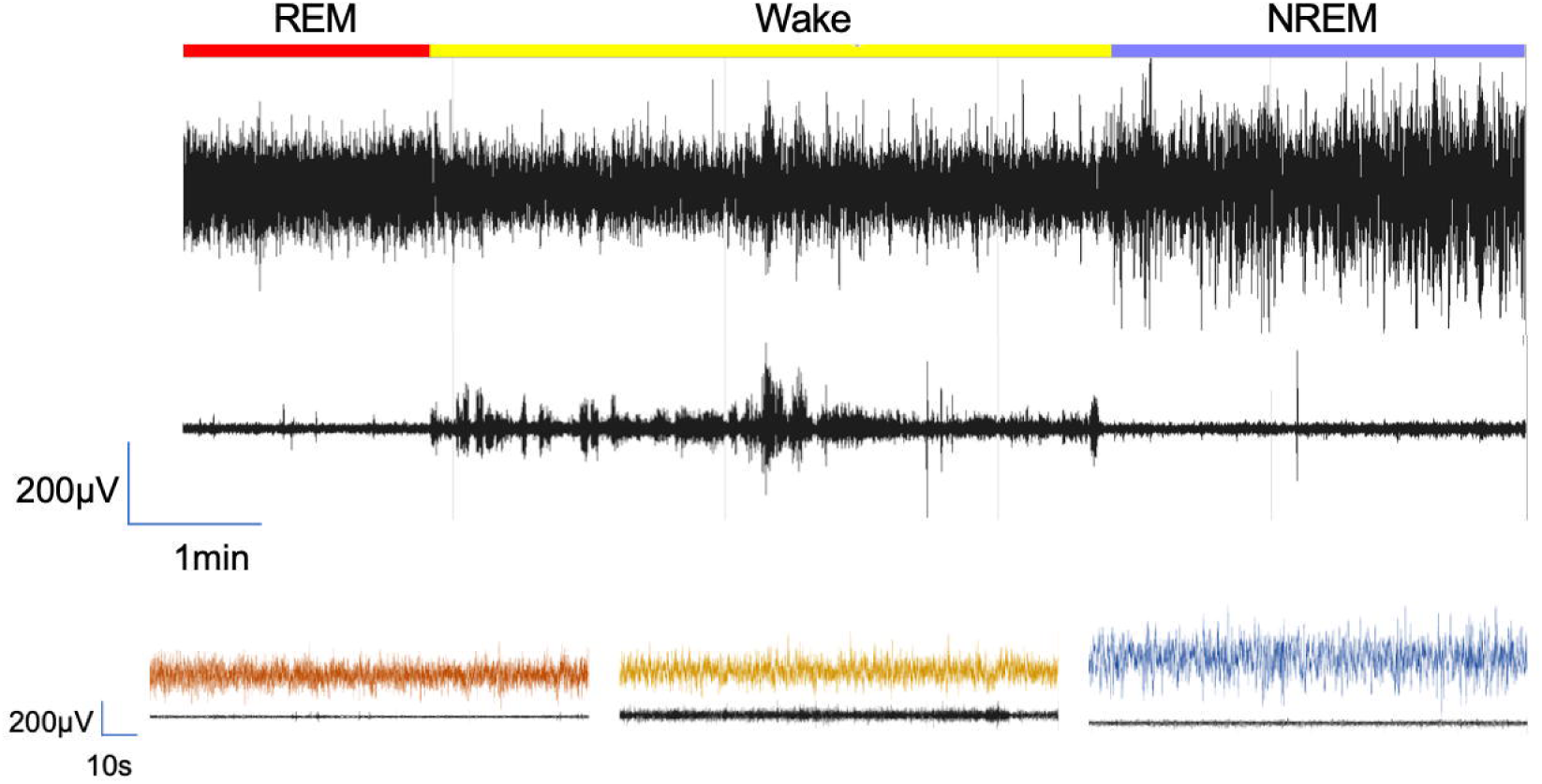
Vigilant state scoring. *Top*: Representative continuous recording of EEG and EMG. *Bottom*: Expanded view of traces from each state. The line colors above the main trace are indicative of the vigilant state represented in the expanded trace.

EEG and EMG data were collected at a sampling rate of 500Hz. The data was acquired by a PC running Dataquest or Ponemah Software (both DSI Inc, St. Paul, Minn.). Digitized EEG and EMG signals were visualized NeuroScore DSI v3.3.9 (DSI Inc, St. Paul, MN) The visualized signals were manually scored in NeuroScore by an experimenter blinded to the treatment groups. The EEG/EMG signals were scored in 10-second epochs. Each epoch was designated as one of three vigilant states as represented in Figure 1: wake (low amplitude, high frequency EEG with high amplitude EMG), NREM sleep (high amplitude, low frequency EEG with low amplitude EMG), or REM sleep (low amplitude, high frequency EEG with muscle atonia or periodic muscle twitches). The total duration (in minutes) for each vigilance state was analyzed for the light and dark phase. Additionally, where noted bout number and bout length distribution was also analyzed for the light and dark phase.

#### Guide cannula implantations

For experiments that required direct microinjections into the brain, parenchyma guide cannulas were implanted. For the median preoptic nucleus (MnPO), a single guide cannula (C315G, 26-gauge; Plastics One) was implanted at a 9° angle at the stereotaxic coordinates 0.45mm posterior/ +1.0mm lateral/ 6.5mm ventral relative to bregma as previously described. ^42^ The cannula and EEG leads were secured together with dental cement. Dummy stylets were used to cover and maintain the guide. The skin along the head was sutured around the guide and dummy cap, and the dorsal incision was closed with wound clips. All animals were treated with antibiotic ointment and topical lidocaine as well as carprofen (5 mg/kg) postoperatively and then allowed 7 days to recover before the start of the experiments.

#### Microinfusions into the MnPO

For all infusions, the dummy stylet was removed from the guide cannula and replaced with a 33-gauge microinfusion needle (Plastics One), extending 0.5 mm below the cannula tip. The needle was connected to a Hamilton 1705 RNR 50µl syringe (Hamilton, Reno, NV) via polyethylene tubing. Infusions were delivered at a rate of 0.1 µl/min (10 mins for the 1µl volume) using a BASi Bee pump and Bee Hive controller (Bioanalytical Systems, Inc., West Lafayette, IN). After each infusion, the needle remained in place for 5 minutes to allow for diffusion.

#### Confirmation of Cannula Placement

At the end of each microinfusion experiment, animals were transcardially perfused with 0.9% saline + 2% sodium nitrite followed by 4% paraformaldehyde in 0.05M KPBS (Potassium Buffered Saline). The brains were removed and post-fixed overnight in 4% paraformaldehyde. Brains were cryoprotected in 30% sucrose in KPBS, frozen on dry ice, and stored at −80°C. Each brain was cut on a cryostat along the coronal plane at 30μm thick into 4 series and stored in an ethylene glycol-based cryoprotectant −20°C. Sections in each series are separated by 120μm.

Sections corresponding to the MnPO were mounted on 2% gelatin-coated slides. The slides were processed for cresyl violet (0.1% solution; cresyl violet acetate, Sigma-Aldrich) staining to examine cannula placement. Cannula placement was determined by visual identification under a light microscope of the guide cannula bore and/or lesion created by cannula insertion. Such points falling within the preoptic area boundary of bregma −0.3mm to bregma +0.4mm and within 1mm above the MnPO (expected tip of the guide cannula) were counted as hits. Supplemental Figures 6A and 9 have representative maps of the injection sites.

### Experiments

#### E2 replacement paradigm overview

In all experiments involving systemic E2 administration, we used our previously validated dosing schedule that mimics the natural peri-ovulatory rise in E2 in cycling female rats and reproduces their sleep–wake patterns.^28,33,38,40,45^ Figure 2A provides a schematic of the standard experimental paradigm. On Day 1 OVX animals receive a subcutaneous (s.c.) injection of sesame oil (50µl; Sigma-Aldrich, St. Louis, MO) ∼15 minutes prior to the start of the light phase (ZT0) for a baseline EEG/EMG. On Day 2, 24 hours after the vehicle injection, the animals receive 5µg E2 (50µl) followed by 10µg (100µl) E2 24 hours later on Day 3. The E2 is dissolved in sesame oil at concentration of 10µg/0.1ml and injected s.c. with a 25Gx 5/8” BD Precision Glide Needle attached to a 1ml TB syringe (Becton, Dickerson and Company, Franklin Lake, NJ). EEG/EMGs are recorded continuously through Day 4. This approach produces circulating E2 levels that peak at physiological proestrus levels following the second E2 injection^45^ and ensures standardized and reproducible hormone concentrations on designated recording days. Our previous results have shown Days 3 and 4 following the last E2 injection are when the most robust changes in sleep wake behavior occur.^38,40,45^ Modifications to this standard E2 replacement paradigm are noted below in the specific experimental designs.

**Figure 2.**
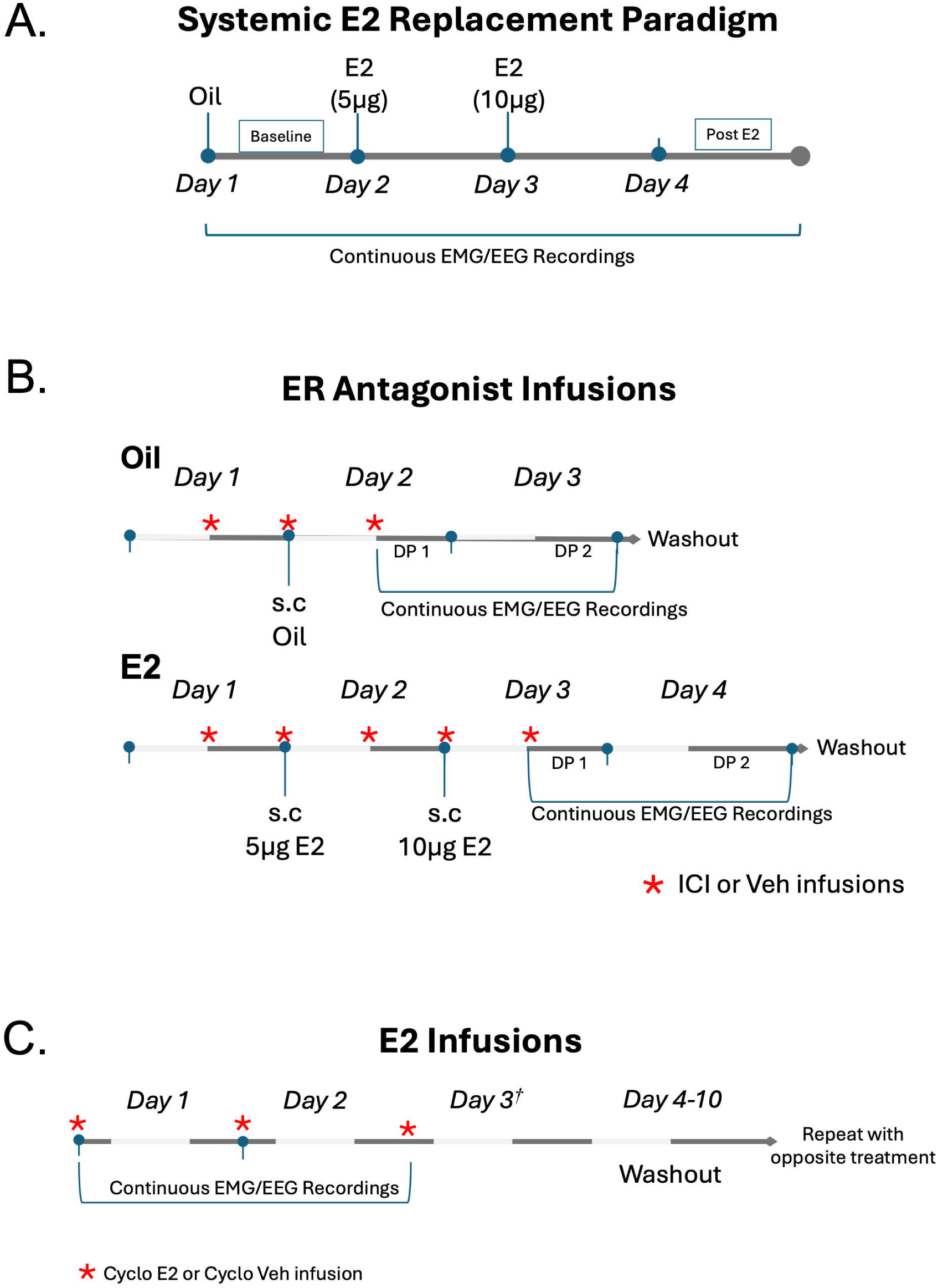
Experimental Paradigms. **(A)** *Schematic of the established E2 replacement paradigm*. This model mimics the natural peri-ovulatory E2 surge and reproduces sleep– wake patterns typically observed in intact females.^40,45^ Ovariectomized (OVX) rats received a subcutaneous oil injection on Day 1, which served as the baseline condition, followed 24 h later by 5 µg E2 (Day 2) and 10 µg E2 (Day 3). Continuous EEG/EMG recordings were collected throughout. This paradigm serves as the basis for the hormonal manipulations used in this study and for the subsequent experimental designs illustrated below. **(B)** *Local infusion of an estrogen receptor antagonist into the MnPO*. To test whether MnPO estrogen receptor signaling is necessary for E2-mediated effects on sleep–wake, the pure ER antagonist ICI 182,780 (Fulvestrant; ICI) or vehicle was infused locally 12 h before, at the time of, and 12 h after the subcutaneous oil or E2 injections. This dosing schedule ensured receptor blockade during the period when exogenous E2 remains elevated.^45^ A within-animal double crossover design was used, consisting of four treatment arms (oil + vehicle, oil + ICI, E2 + vehicle, E2 + ICI). Each arm was followed by a 5-7 day washout before animals received an alternate combination. EEG/EMG recordings were collected for 36 h following each treatment arm. *DP1; Dark phase 1, DP2: Dark phase 2*. **(C)** *Local infusion of estradiol into the MnPO*. To test whether MnPO-specific E2 signaling is sufficient to alter sleep–wake behavior, cyclodextrin-encapsulated E2 (Cyclo-E2) or vehicle (Cyclo-Veh) was infused every 25 hours for three consecutive days at ZT17. Continuous EEG/EMG recordings were started after the first injection. A within-animal crossover design was used with a 7-day washout between treatments. The *†* on Day 3 indicates that data from this time point were incomplete and not included in the analysis.

#### Experiment 1: Effects of Exogenous Progesterone in the Presence of E2

We used the standard exogenous E2 replacement paradigm described above to investigate whether progesterone, in the presence of E2, further influences sleep–wake behavior. Treatment began 14 days after OVX/telemeter implant, with all steroid or vehicle injections administered subcutaneously ∼15 minutes before ZT0. EEG/EMG were continuously recorded starting after the first injection. After completing the E2 replacement protocol outlined above (Figure 2A), animals were randomly assigned to receive either vehicle (Post-E2; 100 µL; n=10) or progesterone (Progesterone; 500 µg/100 µL; n=10) on Day 4. The progesterone dose was selected based on our prior work demonstrating that it produces physiologically relevant levels sufficient to elicit reproductive behaviors in females.^46–48^ Three animals were excluded from analysis due to transmitter loss.

#### Experiment 2: Effects of Specific Estrogen Receptor Agonists on Sleep-Wake Behavior

To investigate the contributions of the classical estrogen receptor subtypes, ERα and ERβ, in sleep-wake behavior, we administered selective agonists for each subtype. Propyl pyrazoletriol (PPT), is a highly selective agonist with approximately 410-fold greater affinity for ERα than ERβ.^49^ Diarylpropiolnitrile (DPN) is an ERβ agonist with a approximately 70-fold greater affinity for ERβ over ERα.^50^ Both compounds were obtained from Tocris, (Ellisville, MO). Each agonist (Tocris, Ellisville, MO) was dissolved separately in DMSO and then diluted in an oil vehicle (10% DMSO:oil). Ovariectomized rats were assigned to one of five treatment groups: oil vehicle (n=8), E2 (as a positive control; n=7), PPT (n=7), DPN (n=7), or a combination of PPT and DPN (n=6).

All treatments were given subcutaneously approximately 15 minutes before ZT0. E2 was administered according to our standard dosing paradigm. Each selective agonist, PPT and DPN, was administered in a low (100 µg/50 µl) dose followed 24 hours later by a high (200 µg/100 µl) dose, mirroring the timing and stepwise increase used in the E2 replacement paradigm (Figure 2A). In the combined selective agonist group, PPT/DPN were each given at the same dose used in the corresponding single-compound treatments but mixed into a single injection (50 µl for the low dose, 100 µl for the high dose). Finally, oil vehicle controls were administered as described for the E2 replacement paradigm, with oil volumes and timing matched to those used for both E2 and agonist treatments. Continuous EEG/EMG recordings were started after the first injection and ended 24 hours after the last injection (see Figure 2A). The doses for PPT and DPN were derived from a survey of the published literature and represent concentrations shown to elicit estrogen-dependent behaviors in other experimental contexts.^51–53^

#### Experiment 3a: Estrogen Receptor alpha (ERα) Immunoreactivity in median preoptic area nucleus and ventrolateral preoptic area

To quantify the ERα positive cells in the MnPO, OVX females (n=8) and sham ORDX males (n=6) were collected during the dark phase. Females received either oil (n=4) or E2 (n=4) according to our standard E2 injection paradigm (Figure 2A), and males received oil (n=6). Twenty-four hours after the last injection animals were overdosed with a ketamine/acepromazine and transcardially perfused with 0.1M Potassium Phosphate Buffered Saline (KPBS) followed by 4% paraformaldehyde in KPBS. Brains were removed, cryoprotected in 30% sucrose in KPBS, frozen on dry ice and stored at −80°C before being sectioned coronally at 30 μm. Sections were stored in ethylene glycol-based cryoprotectant (1% polyvinylpyrrolidone, 30% sucrose, 30% ethylene glycol in PBS, pH 7.2) at −20 °C until processed for ERα immunohistochemistry. Only ERα was examined due to the lack of a reliable ERβ antibody at the time.

All treatment groups were processed together. Sections were rinsed in KPBS, incubated in 1.0% hydrogen peroxide for 30 min to quench endogenous peroxidase, rinsed again, and incubated for 48 h at 4 °C with rabbit polyclonal anti-ERα (cat#06-935, EMD Millipore, Burlington, MA; 1:50,000) in 5% normal goat serum and 0.05% Triton X-100 in KPBS. After rinsing, sections were incubated for 1 h with biotinylated goat anti-rabbit secondary antibodies (1:1000; BA-1000, Vector Laboratories, Burlingame, CA), followed by an avidin–biotin horseradish peroxidase complex (Vectastain ABC Elite Kit; Vector) for 1 h at room temperature. Visualization was achieved with nickel sulfate (25 mg/mL) and diaminobenzidine (0.2 mg/mL; Polysciences, Warrington, PA) in TBS containing 0.005% H₂O₂ for 15 min. Sections were rinsed, transferred to KPBS, mounted on 2% gelatin-coated slides, and coverslipped.

All analyses were performed with the investigator blinded to the experimental groups. For MnPO quantification, anatomically matched, coded slides were analyzed using the optical fractionator workflow in StereoInvestigator (MicroBrightField, Colchester, VT). A two-in-four series (adjacent sections were separated by 60μm; Plates 33–35 of Paxinos and Watson rat brain atlas^54^) was used, with 5–6 sections analyzed per animal. A 100 × 100 μm grid was placed within MnPO contours, and ERα-positive cells were counted within 50 × 50 μm frames. Total estimated cell populations were calculated using the mean section thickness for each animal.

In the VLPO, ERα labeling was too sparse for optical fractionator methods. Instead, all ERα-positive cells were counted in four sections from a one-in-four series (adjacent sections were separated by 120μm; Plates 33 and 36 of the Paxinos and Watson atlas^54^). The placement and size of the VLPO contours were in accordance with previously defined parameters, extending ∼1 mm from the ipsilateral ventricular wall, 0.6 mm laterally, and 0.3 mm dorsally.^55^ Three contours per bilateral section were counted, and an average count per section was calculated.

#### Experiment 3b RNAScope for ESR1 and ESR2 Expression in the MnPO

ERα and ERβ proteins are encoded by the *ESR1* and *ESR2* genes, respectively. In the absence of a reliable ERβ antibody, we examined *ESR1* and *ESR2* mRNA expression in the MnPO to determine whether transcripts for these receptors are present in this region. Brains from OVX females treated with oil (n=3) or E2 (n=3) according to the paradigm described above, and sham-ORDX males treated with oil (n=3), were processed for fluorescent in situ hybridization using the RNAScope system (ACD Bio, Newark, CA). Tissue collection, perfusion, fixation, cryoprotection, freezing, and coronal sectioning at 30 μm followed the procedures described in Experiment 3a. Sections were stored in cryoprotectant until processing.

One day prior to processing, two to three anatomically matched brain sections per animal were selected and mounted onto Fisher SuperFrost Plus slides (Waltham, MA), and air-dried overnight. The selected sections corresponded to anatomical levels represented in Plates 33 to 36 of the Paxinos and Watson rat brain atlas^54^ as described above. Serial sections were spaced approximately 60μm apart to ensure consistent sampling across the region of interest.

*In situ* detection of *ESR1* and *ESR2* mRNA followed the manufacturer’s protocol (ACD Bio). Akoya Opal fluorophores (Marlborough, MA) were used to label probes: *ESR1* with Opal 690, ESR2 with Opal 620. Nuclei were counterstained with DAPI (1:2000 in PBS). Z-stacks from at least three regions of interest (ROIs; 360 × 280 μm) per MnPO were imaged at 40× magnification on a Keyence BZ-X810 microscope, with exposure times and illumination kept constant for each target.

Z-stacks were analyzed in Imaris (Bitplane) following published methods. DAPI-stained nuclei were defined as “Surfaces” (≤10 μm diameter) and *ESR1/ESR2* mRNA signals as “Spots” (>1 μm) in pseudo-colored red (ERα) and green (ERβ) channels, respectively. A cell was considered ERα+ or ERβ+ if ≥3 spots were within 6 μm of the DAPI-defined surface. Binary voxel masking (1 = inside, 0 = outside) refined signal quantification. For each ROI, the percentage of DAPI+ cells containing ERα, ERβ, or both was calculated, averaged across sections for each animal, and then averaged by sex. All image analyses were performed with the investigator blinded to the experimental groups. Representative images are shown in Figure 6B.

#### Experiment 4: Local Infusion of Estrogen Receptor Antagonist into the MnPO

To test whether MnPO estrogen receptor signaling is necessary for systemic E2 effects, ovariectomized (OVX) rats (n = 22) underwent the standard oil/E2 replacement paradigm (Figure 1B) combined with local infusions of the pure ER antagonist ICI 182,780 (ICI; 50 ng in 1 µl sterile saline containing 0.25% DMSO) or vehicle. ICI is a pure antiestrogen without agonist activity.^56–58^ The dose was based on intracerebroventricular studies showing effective blockade of estrogenic actions and was adjusted for intraparenchymal infusion.^59–63^ Animals were implanted with guide cannulas targeting the MnPO during the same surgery as transmitter implantation and OVX.

A within-animal double crossover design was used, consisting of four treatment arms (Figure 2B; oil + vehicle, oil + ICI, E2 + vehicle, E2 + ICI). Infusions were delivered 12 h before, at the time of, and 12 h after each subcutaneous injection to ensure receptor blockade during the 24 h period of circulating E2 since our previous finding demonstrated elevated E2 levels 24 hours after last E2 injection.^45^ EEG/EMG recordings were collected for 36 h after each treatment arm, with a 5–7 day washout between arms.

Of the 22 animals that began the experiment, 17 completed the full paradigm. At the end of the experiment, brains were collected, processed for Nissl staining, and needle placement verified. Four animals had incorrectly placed cannula and were not included in the analysis (Supplemental Figure 6B) and were excluded from the final analysis. In addition, two animals did not have complete recordings across all arms due to transmitter loss. As a result, the final dataset, includes animals that had at least one complete Oil/E2 recordings within the ICI and Vehicle treatment arms (Veh: n=13; ICI: n=11).

In a separate experiment, the role of MnPO ER signaling in males was examined. Intact males (n=8) were implanted with transmitters and guide cannulas targeting the MnPO. Fourteen days post-surgery, animals received a Veh infusion followed by an ICI infusion 25 h later, with EEG/EMG recordings collected for the 24 h immediately following each infusion. After completing the first arm, all males were ORDX, allowed to recover for 14 days, and then treated as in the first arm.

#### Experiment 5: Local Infusion of Estradiol into the MnPO

To test whether E2 signaling within the MnPO is sufficient to alter sleep–wake patterns, we locally infused a water-soluble form of E2 into the MnPO. This consisted of cyclodextrin-encapsulated 17β-estradiol (Cyclo-E2; 5µg total weight; 5% E2, 95% 2-hydroxypropyl-β-cyclodextrin; Sigma-Aldrich) dissolved in 5µl sterile saline. Vehicle (Cyclo-Veh) contained 5µg hydroxypropyl-β-cyclodextrin alone in 5µl sterile saline. The dose was adapted from prior studies showing effective modulation of neural and behavioral outcomes following local Cyclo-E2 administration to discrete brain regions.^64–67^ OVX female rats (n=10) were implanted with transmitters and guide cannulas targeting the MnPO. A within-animal design was used with animals randomly assigned to receive either Cyclo-E2 or Cyclo-Veh in the first treatment arm (Figure 2C).

Infusions were administered at ZT 17 on three consecutive days, 24 h apart, at a rate of 0.3 µl/min, with the infusion needle left in place for 5 min to allow diffusion. EEG/EMG recordings began at the first infusion. Sleep–wake amounts are expressed as % of total time in each recording period to account for the different durations. After a 7-day washout, animals received the alternate treatment. Cannula placement was histologically verified in 8 of 10 animals (Supplemental Figure 9). Animals with misplaced cannulas were excluded from analysis.

The three-day infusion schedule mirrored our systemic E2 replacement paradigm and was selected to capture both rapid and delayed effects of E2. Our previous work shows that many E2-induced changes in sleep emerge ∼24 h after administration, consistent with genomic mechanisms, but Cyclo-E2 has also been reported to produce rapid effects within hours. Infusions at ZT 17 were therefore timed to allow assessment of the immediate response in the remaining dark phase as well as the subsequent light phase. This design allowed us to evaluate both timeframes, with the primary comparison planned for the full 24-h recording on Day 3. Due to a power outage, Day 3 data from one cohort were incomplete, preventing comparison between the treatment groups. Nevertheless, intact Day 1 and Day 2 data provided sufficient information to evaluate the effects of MnPO-targeted E2 infusion on sleep–wake patterns.

### Statistical Analysis

All statistical tests were conducted using the Graph Pad Prism v10 (San Diego, CA). For repeated measure ANOVAs, sphericity was not assumed and a Geisser-Greenhouse’s correction was applied. The specific statistical tests and values are described in detail in the figure legends.

## Results

### Estradiol predominantly influences sleep-wake behavior in the adult female rat during the dark phase

EEG/EMG data from OVX adult rats were assessed across the 3-day steroid treatment paradigm for E2 effects (Figure 2A). E2 administration significantly altered sleep-wake durations primarily in the dark phase (Figure 3A). E2 increased wake and decreased both NREM and REM sleep in a dose-dependent manner, with the most pronounced effects observed on Day 3 following the high dose (10 µg). Post-hoc analysis of hourly totals revealed significant time-specific effects, at ZT13, ZT14, and ZT15, where wake increased on Day 3 compared to baseline. Correspondingly, NREM decreased at ZT13 ZT14, and ZT15, whereas REM was significantly decreased on ZT14, and ZT15 (Figure 3A, *left panel*). Comparison of 12-hour totals further supported an overall effect of E2 in the dark phase on Day 3, where wake significantly increased by approximately 50 minutes with a concomitant decrease in NREM and REM sleep by approximately 35 and 11 minutes, respectively compared to baseline (Figure 3A, *right panel*).

**Figure 3.**
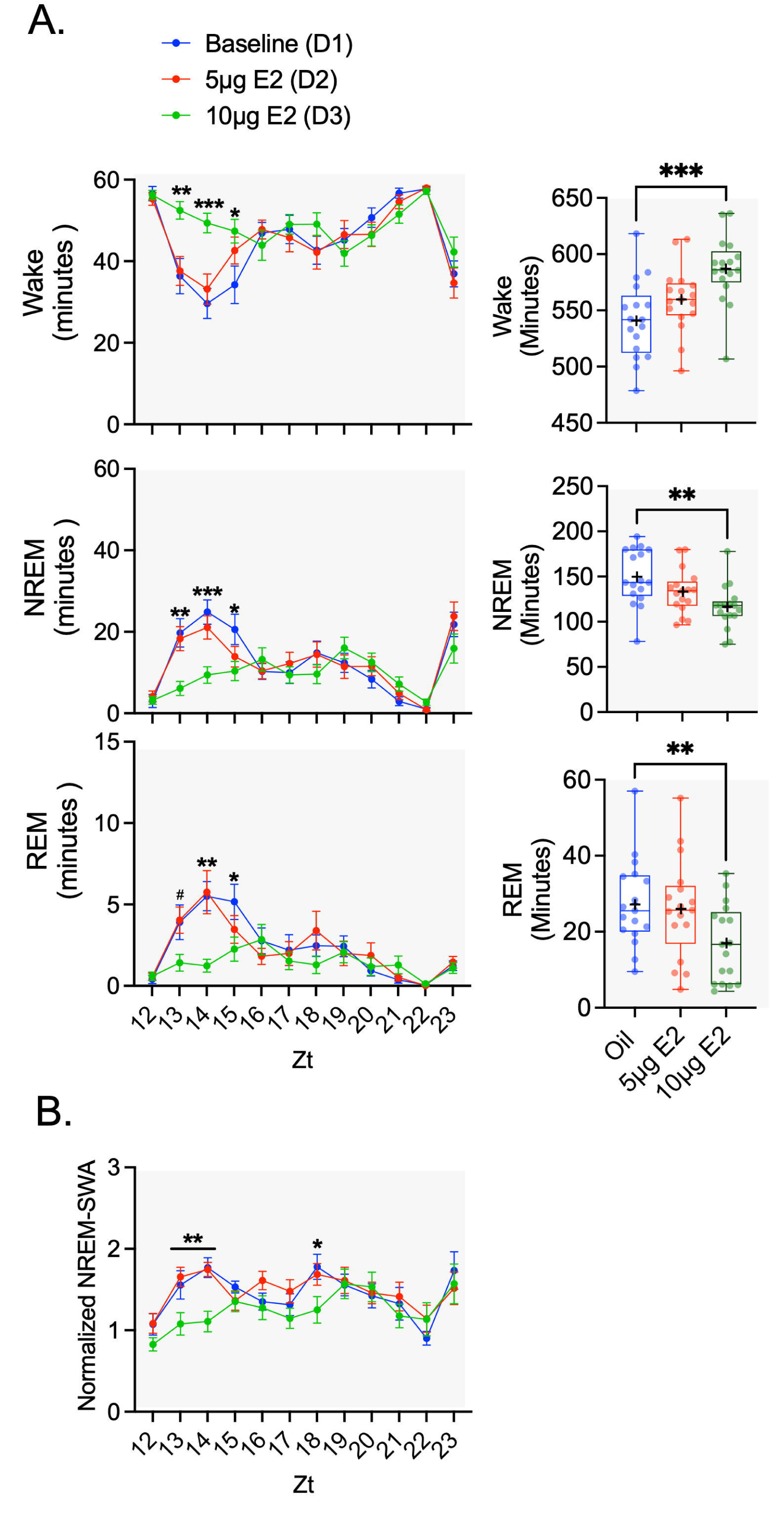
Estradiol predominantly influenced sleep-wake behavior in adult female rats during the dark phase. Ovariectomized (OVX) rats received systemic oil and E2 injections across three consecutive days according to our standard paradigm (Figure 2A). **(A)** *Left*, Hourly means of wake, NREM, and REM during the dark phase. Systemic E2 significantly increased wake and decreased both NREM and REM, with the largest effects observed on Day 3 (10 µg E2). Time-specific post hoc analysis revealed increased wake and decreased NREM at ZT13, ZT14, and ZT15 compared to oil baseline, while REM was decreased at ZT14 and ZT15, with a trend toward a decrease at ZT13 (#p = 0.06). *Right*, Boxplots of total 12-h dark phase durations. On Day 3, wake increased by ∼50 min, accompanied by decreases of ∼35 min in NREM and ∼11 min in REM compared with oil baseline. **(B)** Hourly means of NREM-SWA during the dark phase. Systemic E2 decreased NREM-SWA, with significant reductions observed at ZT13, ZT14, and ZT18 on Day 3 compared with oil baseline. <u>Statistical analyses:</u> (A) *Left* (hourly means): Two-way repeated measures ANOVA (Treatment × Time, ZT). **Wake**: Main effect of treatment, F_(1.8, 27.9)_=16.46, p<0.0001; Main effect of time, F_(5.2, 83.7)_=12.70, p<0.0001; Interaction, F_(7.7, 122.5)_=2.67, p<0.01. **NREM:** Main effect of treatment, F_(1.8, 29.4)_=13.85, p<0.0001; Main effect of time, F_(5.2, 83.4)_=14.51, p<0.0001; Interaction, F_(7.4, 118.3)_=2.61, p<0.05. **REM:** Main effect of treatment, F_(1.6, 26.2)_=9.21, p<0.0001; Main effect of time, F_(5.1, 80.9)_=8.78, p<0.0001; Interaction, F_(5.8, 93.8)_=2.45, p<0.05. **Post hoc (Dunnett’s):** On Day 3, wake was increased at ZT13 (**p<0.005), ZT14 (***p< 0.0005), and ZT15 (*p<0.05) compared to oil baseline. NREM was decreased at ZT13 (**p<0.005), ZT14 (***p<0.001), and ZT15 (*p<0.05) compared to oil baseline. REM was decreased at ZT14 (***p< 0.0005) and ZT15 (*p< 0.05), with a trend at ZT13 (#p =0.06) compared to oil baseline. (A) *Right*, (12-h dark phase totals): One-way repeated measures ANOVA. Wake: F_(1.5, 24)_=15.53, p<0.0001 NREM: F_(1.4, 22.9)_=13.9, p<0.0005 REM: F_(1.7, 26.3)_= 8.7, p<0.005 Post hoc (Dunnett’s): On Day 3, wake was significantly increased (**p<0.001) and both NREM and REM were significantly decreased (**p<0.005) compared to oil baseline. (B) (NREM-SWA hourly means): Mixed-effects model (Treatment × Time, ZT). Main effect of treatment: F_(1.163, 18.61)_=4.369, p<0.05 Main effect of time: F_(3.27, 52.36)_=4.93, p<0.01 Post hoc (Tukey’s): NREM-SWA was reduced on Day 3 at ZT13 (**p<0.001), ZT14 (**p<0.01), and ZT18 (*p<0.05). All p values were adjusted for multiple comparisons. For box plots, all data points are shown; the line represents the median and the + represents the mean.

Light-phase effects are provided in Supplemental Figure 1A. There was little to no differences in hourly totals across the light phase for each state. The 12-hour totals revealed modest changes in sleep–wake behavior. Compared to oil control, overall time spent in wake on Day 3 decreased by approximately 20 minutes following E2 treatment with similar concomitant increase in NREM sleep. REM sleep was unchanged.

### Estradiol decreases NREM-SWA in the adult female rat during the dark phase

NREM-SWA, defined as the amount of delta (0–4 Hz) power in EEG signals during NREM sleep bouts, is a key marker of sleep homeostasis. Analysis revealed that E2 administration significantly decreased NREM-SWA in the dark phase (Figure 2B). Hourly analyses demonstrated significant decreases at ZT13, ZT14, and ZT18, on Day 3 following the high dose of E2 (10 µg), compared to baseline. In the light phase, hourly NREM-SWA levels showed a clear time-of-day pattern but no effect of E2 treatment (Supplemental Figure 1B).

### Progesterone in the presence of E2 has no significant effect on sleep-wake states compared to E2-alone

The rat estrous cycle is characterized by a sharp rise in progesterone following the peak of E2 on proestrus. Typical for our model, the effects of E2 persist 24 hours after the final injection (D4; Figure 2A).^40^ Additionally, we have shown circulating E2 levels remain elevated at this time point.^45^ Together, this presents an opportunity to mimic the natural proestrus progesterone surge and assess whether progesterone contributes to the effects of E2 on sleep–wake behavior. To test whether this rise in progesterone further influences sleep-wake behavior, the E2-treated animals (above) were divided into two groups: Post-E2 and E2 + Progesterone. On D4, 24 hours after the final E2 treatment, animals received either a physiological dose of progesterone (500 mg) or an oil vehicle.

First, to confirm the persistence of E2 action in the present study, sleep–wake durations in the Post-E2 group were compared across baseline (D1), the day of the last injection (D3), and the following day (D4). Consistent with previous findings, increased wake and decreased NREM and REM sleep during the dark phase on D4 were sustained, and D4 values were not significantly different from those observed on D3 (Supplemental Figure 2). Next, sleep-wake durations in the light and dark for the Post-E2 and E2 + Progesterone groups were compared to baseline (D1). No significant differences in wake, NREM, or REM sleep were observed between the Post-E2 and E2+P groups in either the light or dark phases. Compared to baseline Day 1, both groups exhibited significant increases in wake and decreases in NREM and REM sleep durations in the dark phase (Supplemental Figure 3).

### Estradiol consolidates sleep–wake architecture in the first half of the dark phase

Rodents are polyphasic sleepers and exhibit numerous sleep–wake transitions throughout both the light and dark phases. Because E2 altered overall sleep–wake amounts, we next asked whether these effects were accompanied by changes in sleep architecture. Bout analysis in 6 h bins on Day 3 compared to Day 1 revealed that E2 reduced the number of wake, NREM, and REM bouts in the first half of the dark phase (ZT12–17), but not in the second half (ZT18–23) (Figure 4A). During ZT12–17, E2 also increased mean bout duration for wake while decreasing the bout duration for NREM and REM (Figure 4B). These data suggest that E2 consolidated wake in the dark phase by producing fewer and longer wake bouts at the expense of fewer and shorter NREM and REM bouts. In contrast, effects in the light phase were modest. E2 reduced wake and NREM bout number without altering bout duration (Supplemental Figure 4A,B).

**Figure 4:**
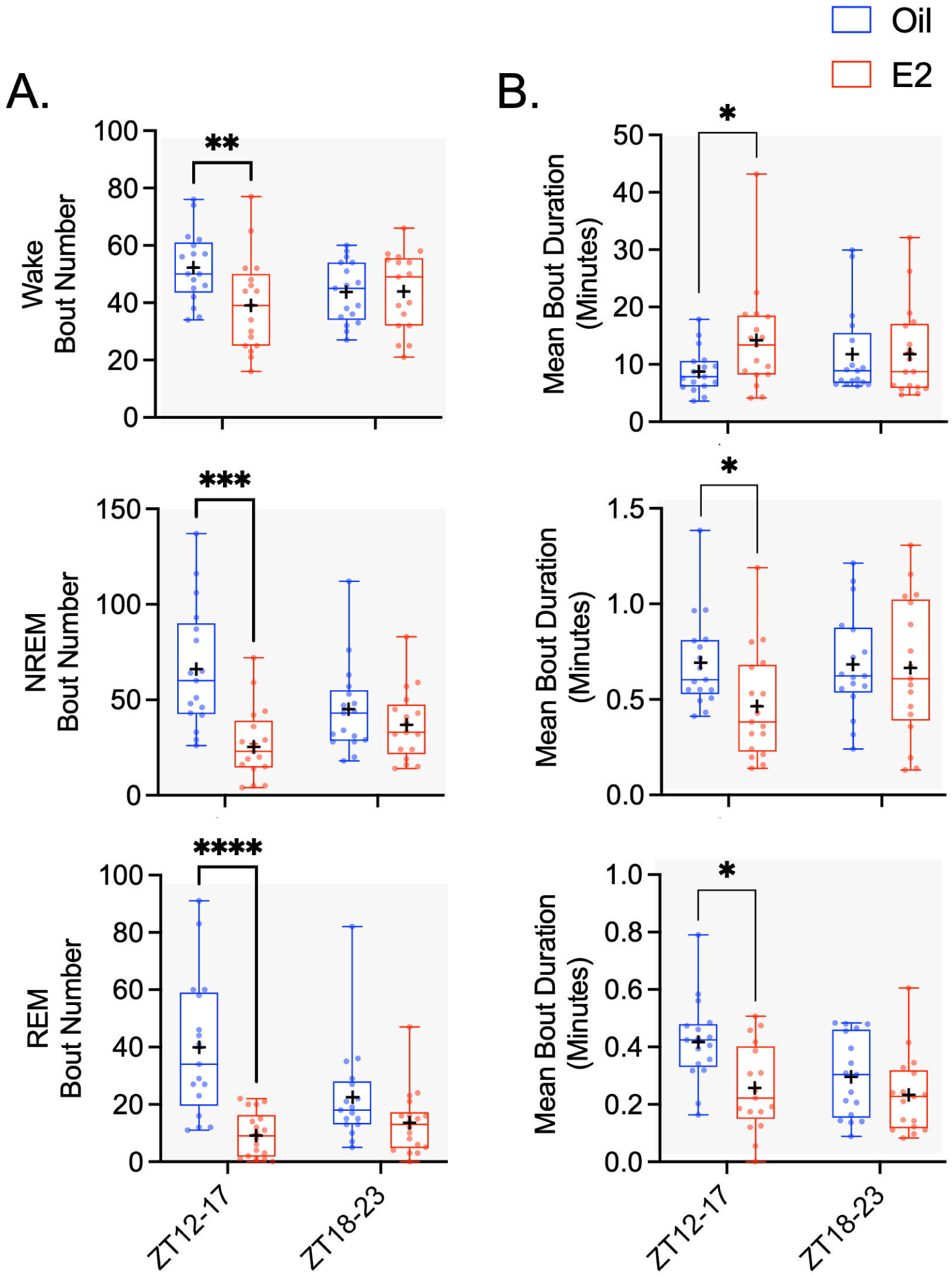
Estradiol consolidates sleep–wake architecture in the first half of the dark phase. Sleep architecture was assessed by comparing bout number and mean bout duration in 6 h bins during the dark phase at baseline (Day 1) and following E2 treatment (Day 3, see Figure 2A). **(A)** E2 reduced the number of wake, NREM, and REM bouts during the early dark phase (ZT12–17), but no changes were observed in the late dark phase (ZT18–23). **(B)** E2 increased the mean bout duration of wake while decreasing the length of NREM, and REM bouts during the early dark phase (ZT12–17). No changes were observed in the late dark phase (ZT18–23). <u>Statistical analyses:</u> (A) two-way repeated measures ANOVA (Treatment × Time): **Wake:** Interaction, F_(1,16)_=11.7, p<0.005; trend for a main effect of treatment, F_(1,16)_=4.0, p=0.05. **NREM:** Main effect of treatment, F_(1,16)_=19.3, p<0.001; interaction, F_(1,16)_=29.9, p<0.0001. **REM:** Main effect of treatment, F_(1,16)_=8.5, p<0.01; main effect of time, F_(1,16)_=31.8, p<0.0001; interaction, F_(1,16)_=22.4, p<0.0005. **Post hoc (Šídák’s):** During ZT12–17, bout number was reduced on D3 compared to baseline for wake (**p<0.01), NREM (***p<0.0005), and REM (****p<0.0001). (B) two-way repeated measures ANOVA (treatment × time): **Wake:** Interaction, F_(1,16)_=5.1, p<0.05. **NREM:** Main effect of treatment, F_(1,16)_=5.2, p<0.05. **REM:** Main effect of treatment, F_(1,16)_=7.3, p<0.05; interaction, F_(1,16)_=4.9, p<0.05. **Post hoc (Šídák’s):** During ZT12–17, mean bout duration was increased for wake (*p<0.05) and decreased for NREM (**p<0.005) and REM (*p<0.05) on D3 compared to baseline. All p values were adjusted for multiple comparisons. Data are presented in box plots with all data points; the line represents the median and the + represents the mean.

### Activation of both estrogen receptor alpha and estrogen receptor beta receptors are necessary for changes sleep-wake behavior

ERα, and ERβ are the primary nuclear receptor subtypes in the CNS, but their individual roles in regulating sleep-wake behavior remain unclear. Although both receptors are widely distributed, their individual contributions to sleep–wake regulation remain unclear. To address this, we tested the effects of selective ERα and ERβ agonists, alone and in combination, to determine whether activation of either receptor subtype was sufficient to replicate the sleep–wake effects of E2. OVX rats were treated with PPT (agonist for ERα), DPN (agonist for ERβ), PPT+DPN, E2, or vehicle. All agonists were administered in a stepwise fashion, with a low dose on Day 1, a high dose on Day 2, and recordings continued through Day 3, the post-injection period 24 h after the final treatment (Figure 2A). This dosing schedule was designed to mimic the gradual rise in estradiol and match our established replacement paradigm. Because E2 effects were most pronounced in the dark phase, analysis of sleep–wake durations was limited to this period.

Figure 5 shows the comparisons of dark phase Wake, NREM and REM durations across all five treatment groups on the post-injection day (D3). At this time point, E2 and the combined PPT+DPN treatment significantly increased wake and decreased both NREM and REM sleep compared to oil, whereas PPT or DPN alone were not significantly different from the oil control. Supplemental Figure 5 shows sleep–wake durations across the low-dose (D1), high-dose (D2), and post-injection (D3) days for each treatment group. Across this timeline, only E2 and the combined PPT+DPN treatment produced significant changes between the low-dose and post-injection days, with increased wake and reduced NREM and REM. In contrast, PPT or DPN alone did not differ across days.

**Figure 5:**
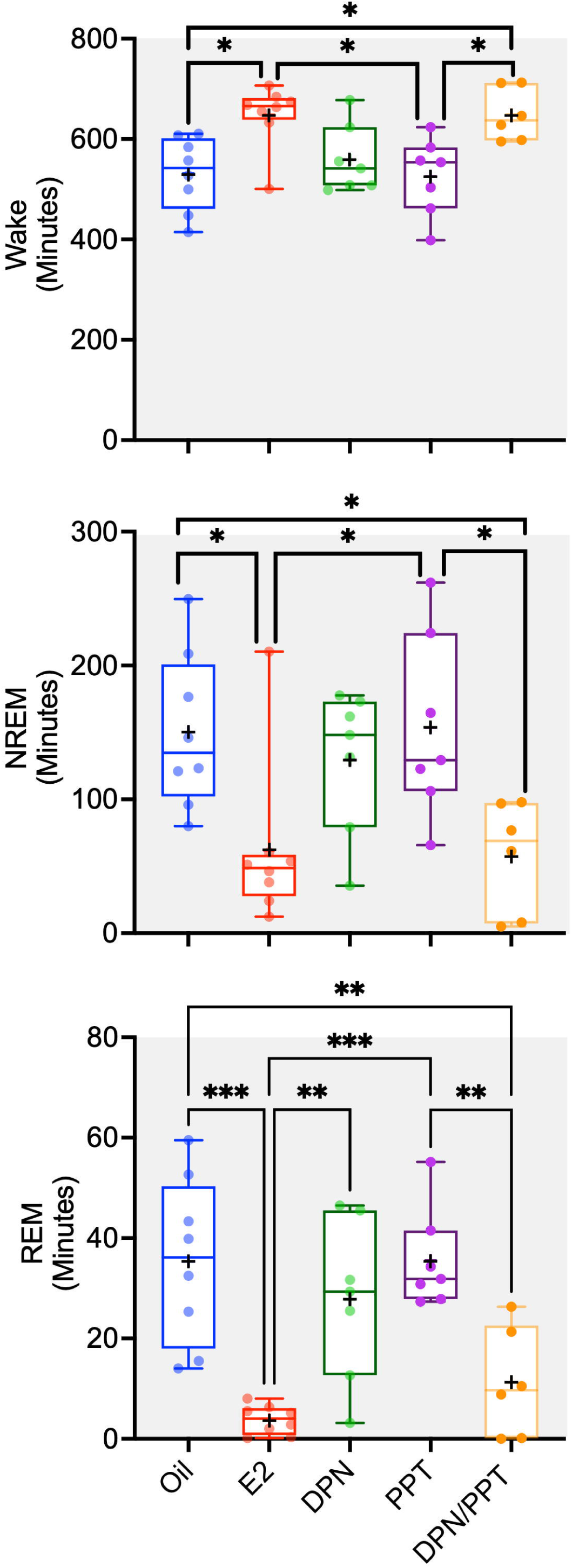
ERα and ERβ are necessary for changes sleep-wake behavior in the dark phase. OVX rats were treated with selective ERα (PPT) or ERβ (DPN) agonists, alone or in combination, along with E2 and vehicle controls. Agonists were administered in a stepwise manner, with a low dose on Day 1 followed by a high dose on Day 2, to mimic the gradual rise in estradiol and match the established replacement paradigm (Figure 2A). Sleep–wake durations were assessed in the dark phase on the post-injection day (D3), 24 h after the final treatment. At this time point, both E2 and PPT+DPN significantly increased wake and decreased NREM and REM compared with vehicle. PPT or DPN alone did not differ from vehicle. Additional group comparisons revealed that E2 and PPT+DPN had significantly more wake and less NREM and REM than PPT, and E2 reduced REM compared with DPN. <u>Statistical analyses:</u> one-way ANOVA with Tukey’s multiple comparisons test: **Wake:** Main effect of treatment, F_(4,31)_=5.9, p<0.005. Post hoc analysis showed that both E2 (p<0.05) and PPT+DPN (p<0.01) animals spent significantly more time awake compared with vehicle. In addition, wake was greater in the E2 (p<0.05) and PPT+DPN (p<0.05) groups compared with PPT. **NREM:** Main effect of treatment, F_(4,31)_=4.7, p<0.005. Post hoc analysis showed that both E2 (p<0.05) and PPT+DPN (p<0.05) groups had significantly less NREM sleep compared with vehicle. NREM sleep was also reduced in the E2 (p<0.05) and PPT+DPN (p<0.05) groups compared with PPT. **REM:** Main effect of treatment, F_(4,31)_=10.4, p<0.0001. Post hoc: Post hoc analysis revealed that both E2 (p<0.01) and PPT+DPN (p<0.001) animals had significantly less REM sleep compared with vehicle. REM was also reduced in the E2 (p<0.05) and PPT+DPN (p<0.05) groups compared with PPT, and E2 animals had less REM sleep compared with DPN (p<0.05). All p values were adjusted for multiple comparisons. Data are presented in box plots with all data points; the line represents the median and the + represents the mean.

### Sex and region-specific differences in ERα and ERβ expression in the MnPO and VLPO

Both *ESR1* and *ESR2* gene expression as well as their protein products (ERα and ERβ), have been identified in the preoptic area nuclei.^68–71^ Additionally, our previous work suggests that both of the sleep associated nuclei, MnPO and VLPO, represent putative sites of E2 action.^42^ To further investigate the discrete localization of estrogen receptors in these nuclei, we quantified protein levels of ERα and mRNA transcripts of *ESR1* and ESR2 in GDX and intact male rats and OVX female rats with and without exogenous E2. Preliminary analysis revealed that the number of ERα-ir cells in the VLPO and MnPO in both males and females was not affected by hormonal status, allowing the data to be combined by sex. The number of ERα -positive (ERα+) cells was significantly higher in the MnPO compared to the VLPO in both male and female rats (Figure 6A). In females, the MnPO contained approximately 30 times more ERα+ cells than the VLPO, while in males, this difference was approximately 19-fold. A sex difference was observed in the MnPO, where females had significantly more ERα-ir cells than males.

**Figure 6:**
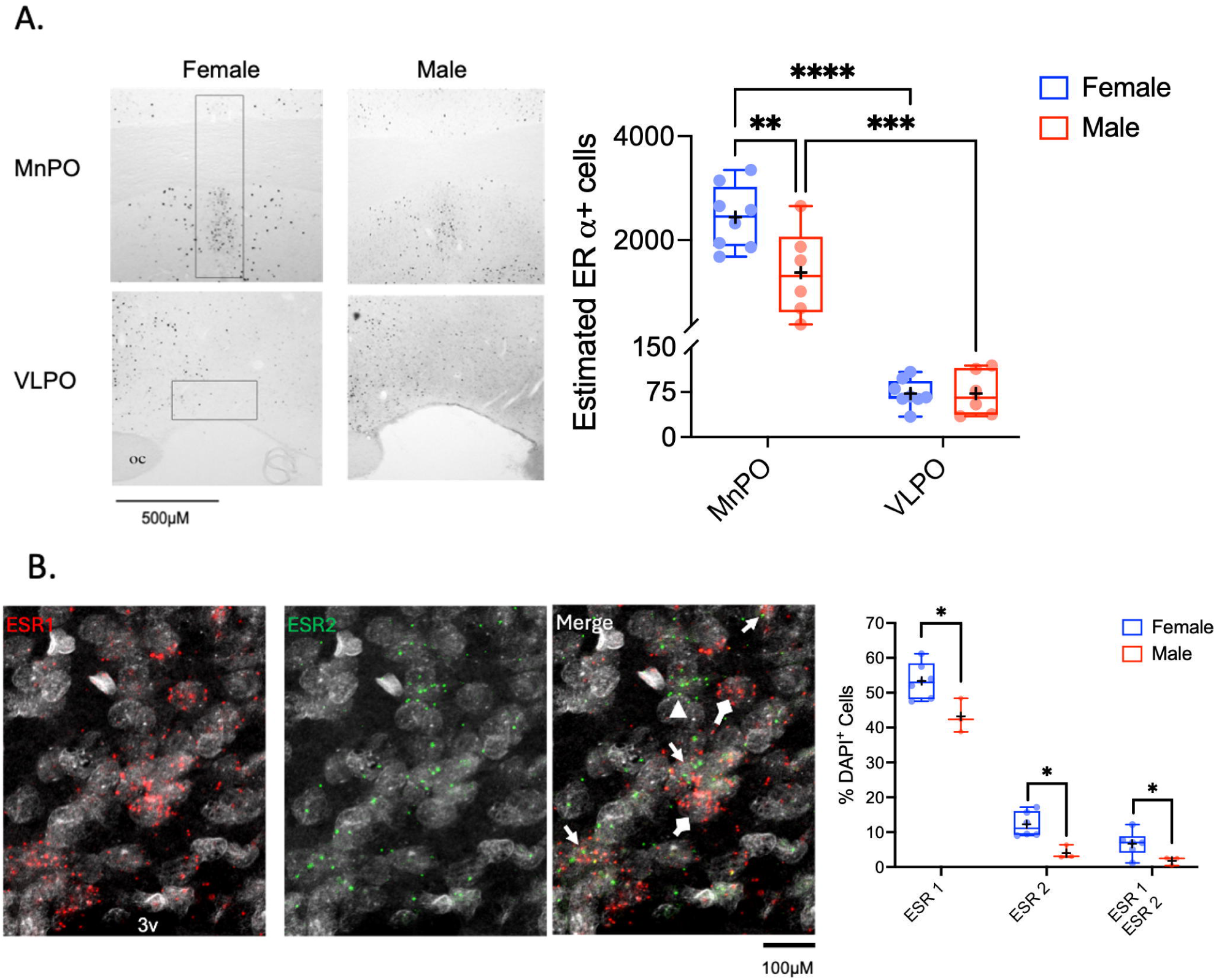
Estrogen receptor localization in the median preoptic nucleus (MnPO) and ventrolateral preoptic area (VLPO) **(A)** *Left*, Representative photomicrographs of ERα immunoreactivity (-ir). *Right*, ERα-ir cells were quantified in the MnPO and VLPO of male and female rats. The number of ERα-ir cells was significantly greater in the MnPO compared with the VLPO in both sexes. In females, the MnPO contained ∼34-fold more ERα-ir cells than the VLPO, whereas in males this difference was ∼19-fold. A sex difference was observed in the MnPO, where females had significantly more ERα-ir cells than males. **(B)** *Left*, Representative photomicrograph series of RNAscope fluorescent in situ hybridization for *ESR1* and *ESR2* mRNA in the MnPO. *ESR1* only cells, block arrow; *ESR2* only cells, arrow heads; co-labeled, point arrow. *Right*, RNAscope fluorescent in situ hybridization was used to quantify *ESR1* and *ESR2* mRNA in the MnPO. Both sexes showed significantly higher numbers of DAPI+ cells expressing *ESR1* compared to *ESR2*. Females had more DAPI+ cells expressing *ESR1* and *ESR2* than males, and co-expression of *ESR1* and *ESR2* was observed in ∼8% of DAPI+ cells in females and ∼2% in males. <u>Statistical analyses:</u> **(A)** two-way repeated measure ANOVA (Region x Sex) with a Šídák’s multiple comparison test. **ERα-ir cells:** Main effect of region, F_(1,20)_=122.3, p<0.0001; Main effect of sex, F_(1,20)_=11.7, p<0.01; interaction, F_(1,20)_=11.7, p<0.01. **Post hoc (Šídák’s):** In females, the MnPO contained significantly more ERα-ir cells than the VLPO (****p<0.0001). In males, the MnPO also contained more ERα-ir cells than the VLPO (***p<0.001). Only the MnPO exhibited a significant sex difference, with females showing higher counts than males (**p<0.01). **(B)** two-way repeated measure ANOVA (Region x Sex) with Tukey’s multiple comparison test. Main effect of receptor isoform, F_(1.03,7.23)_=731.0, p<0.0001 Main effect of sex, F_(1,7)_=8.6, p<0.05; Interaction, F_(2,14)_=3.95, p<0.05. Post hoc analysis (Tukey’s): The percentage of DAPI+ cells expressing ESR1 was significantly higher in females than in males (*p<0.05). The percentage of DAPI+ cells expressing *ESR2* was also greater in females than in males (*p<0.05). In addition, the percentage of co-labeled *ESR1*/*ESR2* cells was higher in females than in males (*p<0.05). All p values were adjusted for multiple comparisons. Data are presented in box plots with all data points; the line represents the median and the + represents the mean.

**Figure 7:**
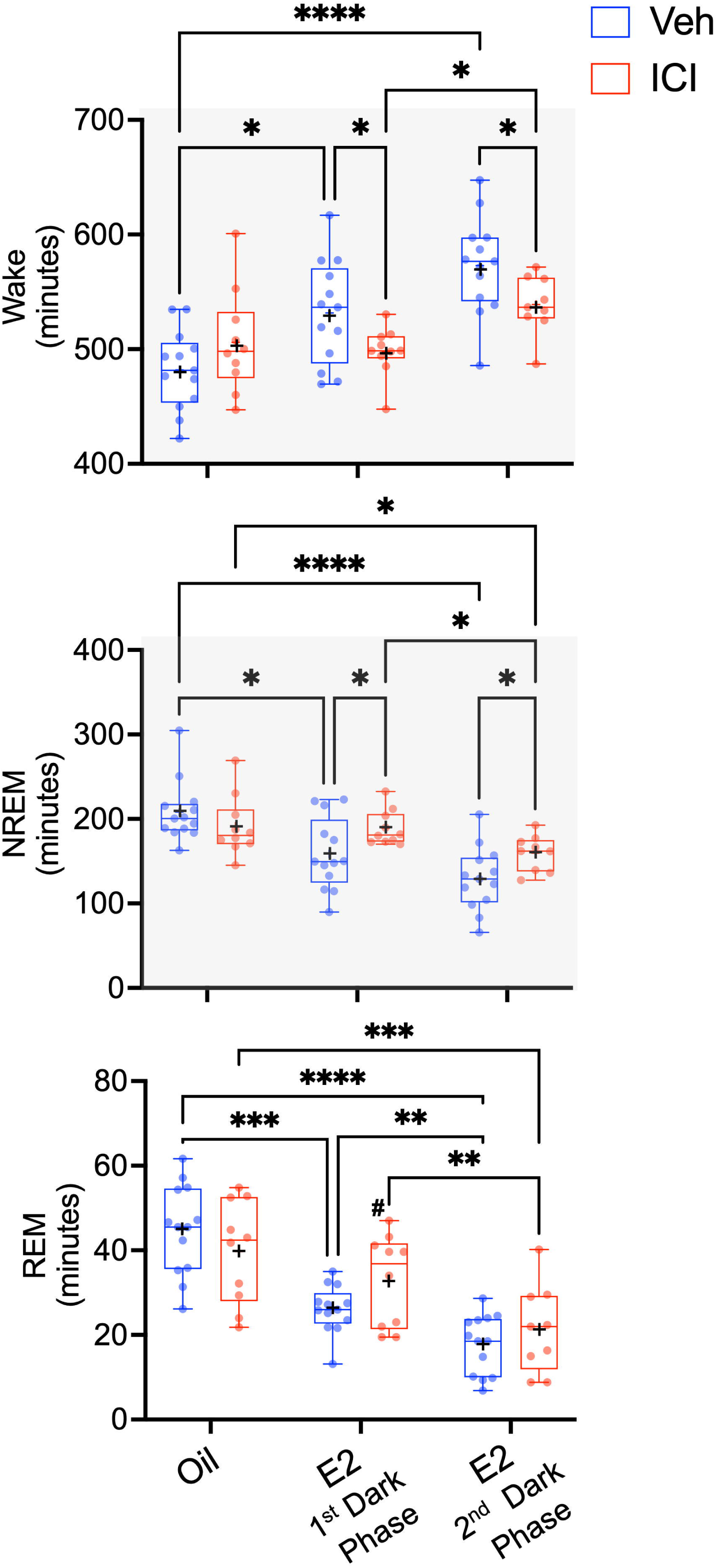
Local infusion of ICI 182,780, directly into the MnPO attenuates E2 effects on sleep-wake states in the dark phase. Ovariectomized (OVX) rats received oil or E2 injections combined with local MnPO infusions of vehicle (Veh) or the estrogen receptor antagonist ICI 182,780 (Fulvestrant; ICI; 50 ng/µl). A within-animal double crossover design was used, generating four treatment arms: Oil/Veh, Oil/ICI, E2/Veh, and E2/ICI (see Figure 2B). Boxplots show total minutes of wake (top), NREM (middle), and REM (bottom) during the oil condition and across both the first and second dark phases following E2 treatment. Systemic E2 (E2/Veh) significantly increased wake and decreased NREM and REM compared with Oil/Veh. ICI had no effect in oil groups (Oil/Veh vs Oil/ICI) but significantly attenuated the actions of E2 for wake and NREM but not REM. E2/ICI animals showed reduced wake and increased NREM compared with their paired E2/Veh controls. REM while not significant showed a tend toward an increase compared to Oil/Veh in the first dark phase (#p=0.07). Data were analyzed using a mixed-effects model. All p-values were adjusted for the multiple comparisons. Data are presented in box plots with all data points. The line represents the median and the **+** represents the mean. <u>Statistical analyses:</u> **Wake:** Main effect of E2: (F_(1.7, 35.4)_=19.60 p<0.0001); ICI×E2 interaction: (F_(1.7, 35.4)_=5.7 p<0.01). **NREM:** Main effect of E2: (F_(1.5, 32.5)_=22.35 p<0.0001); ICI×E2 interaction: (F_(1.5, 32.5)_=5.4, p<0.05). REM: Main effect of E2: (F_(1.5, 29.5)_=58.1, p<0.0001); ICI×E2 interaction: (F_(1.5 29.5)_=4.6, p<0.01). Post hoc comparisons were conducted using Tukey’s multiple comparisons test; *p < 0.05, **p<0.01, ***p<0.001, ****p<0.0001.

To investigate the potential expression of ERβ in the MnPO, we utilized fluorescent in situ hybridization (RNAscope) to localize *ESR1* and *ESR2* mRNA in both female and male rats (Figure 6B). Similar to the cells that expressed ERα, E2 treatment did not significantly affect the percentage of DAPI-labeled cells expressing *ESR1* or *ESR2*; therefore, the treatment groups for females were combined into a single group for analysis. Both male and female rats showed significantly higher numbers of DAPI+ cells expressing *ESR1* compared to *ESR2*, with ∼6.7- and ∼19-fold difference, respectively. A sex difference in the number of DAPI+ cells expressing *ESR1* and *ESR2* alone was detected with females having more DAPI+ cells expressing *ESR1* (1.3 fold) and *ESR2* (4 fold) than males. Co-expression of *ESR1* and *ESR2* was observed in ∼8% of DAPI+ cells in females and 2% in males within the MnPO.

### The ER antagonist, ICI 182,780, infused directly into the MnPO attenuates E2-induced changes in sleep-wake behavior

To test whether MnPO estrogen receptor signaling is necessary for the systemic E2 modulation of sleep–wake behavior, OVX females underwent our standard oil/E2 replacement paradigm combined with local MnPO infusions of the estrogen receptor antagonist ICI 182,780 (Fulvestrant; ICI) or vehicle (Veh). Cannula placements within the MnPO were confirmed histologically, and representative injection sites are shown in Supplemental Figure 6. A within-animal double crossover design generated four treatment arms: Oil/Veh, Oil/ICI, E2/Veh, and E2/ICI (Figure 2B).

In vehicle-infused animals, systemic E2 significantly increased wake and decreased both NREM and REM sleep compared with the oil baseline, replicating our previous findings. The largest effects were observed during the second dark phase (Day 4; 24 h after the high-dose E2 injection). ICI infusions alone had no effect in oil-treated animals, as wake, NREM, and REM durations did not differ between Oil/Veh and Oil/ICI treatments. In contrast, during E2 treatment, ICI attenuated the E2 effect in wake and NREM but not REM. When compared to the E2/Veh treatment, E2/ICI animals showed decreased wake and increased NREM, returning to levels that were not significantly different from oil controls. Interestingly, while E2 effects on REM was not significantly attenuated by ICI, there was a trend toward increased levels in the first dark phase after E2/ICI treatment. In the light phase, systemic E2 had no significant effects on sleep– wake behavior. Wake, NREM, and REM did not differ between E2/Veh and Oil/Veh, and ICI infusions in the presence or absence of E2 did not alter any vigilance state (Supplemental Figure 7).

While ICI significantly attenuated the actions of E2 in Wake and NREM when compared to the paired Veh controls, the effect was not completely reverse. The average percent change across both dark phases following E2 treatment (E2/Veh vs Oil/Veh) was an increase of ∼17% in wake and a reduction of ∼30% in NREM. In comparison, the average percent change between E2/Veh and E2/ICI was ∼8% for wake, and ∼19% for NREM.

In males, a within-animal design was used to test whether endogenous E2, derived from systemic circulation or local de novo synthesis in the brain, influences sleep–wake behavior via MnPO ER signaling. Intact males received vehicle and ICI infusions into the MnPO, followed by EEG/EMG recordings. Animals were then gonadectomized, allowed to recover, and retested with the same infusion protocol. Across all conditions, wake, NREM, and REM did not differ from intact vehicle baselines, indicating that MnPO ER signaling does not significantly contribute to sleep–wake regulation in males (Supplemental Figure 7).

### Estradiol infused directly into the MnPO is sufficient to change sleep-wake behavior

To determine if E2 signaling localized to the MnPO is sufficient to increase wakefulness and suppress NREM and REM sleep, OVX females received local infusions of cyclodextrin-encapsulated estradiol (Cyclo-E2) or vehicle (Cyclo-Veh) in a within-animal design randomized crossover design (Figure 2C). Infusions were delivered once daily for three consecutive days at ZT17, paralleling the systemic replacement paradigm. Sleep–wake amounts were expressed as percentage of total recording time to account for differences in phase duration. Day 2 was the primary focus of analysis, as no significant effects were observed on Day 1. Paired two-tailed t tests were used for all comparisons.

During the first dark period following infusion (DP-1; ZT18–23), Cyclo-E2 significantly increased wake and decreased NREM compared with Cyclo-Veh, whereas REM was unchanged (Figure 8). Similar effects were observed in the subsequent light phase (ZT0–11), where wake was elevated and NREM was reduced. In contrast, by the second dark phase (DP-2; ZT12–17), sleep–wake amounts did not differ between Cyclo- E2 and Cyclo-Veh, indicating that the effects of local MnPO E2 infusion were transient.

**Figure 8:**
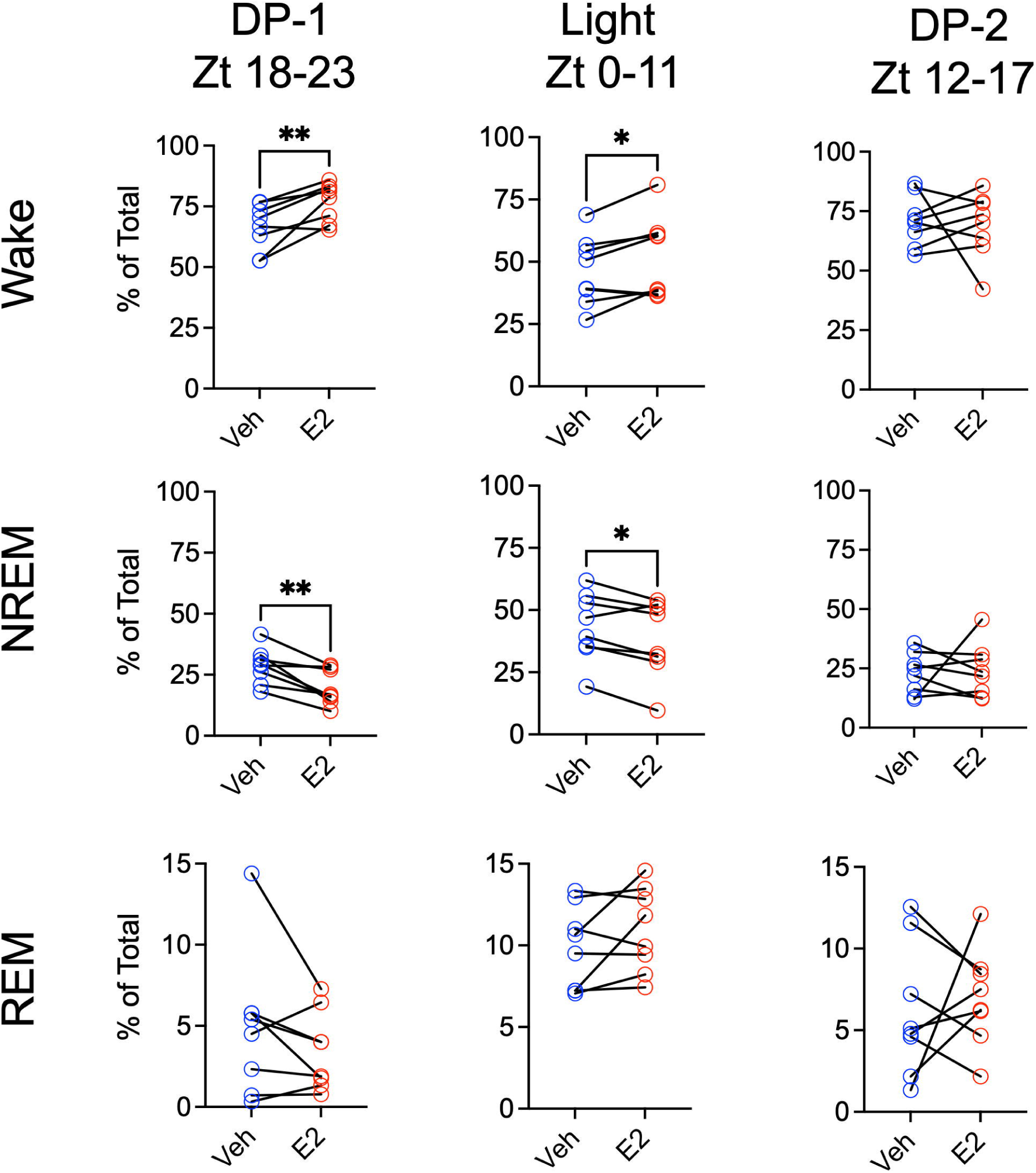
Estradiol infused directly into the MnPO of OVX females is sufficient to change sleep-wake behavior. Ovariectomized (OVX) females received local infusions of cyclodextrin-encapsulated estradiol (Cyclo-E2; 5 µg total) or vehicle (Cyclo-Veh) into the MnPO once daily for three consecutive days at ZT17 (see Figure 2C). Sleep–wake amounts were expressed as percent of total recording time to account for differences in phase duration. Box plots show wake (top), NREM (middle), and REM (bottom) during the first dark phase following infusion (DP-1, ZT18–23), the subsequent light phase (ZT0– 11), and the second dark phase (DP-2, ZT12–17) on Day 2. Cyclo-E2 significantly increased wake and decreased NREM compared with Cyclo-Veh during DP-1 and the light phase, whereas REM was unaffected. By DP-2, sleep–wake durations did not differ between treatments, indicating that the effects of local E2 infusion were transient. <u>Statistical analyses:</u> paired two-tailed t tests: **Wake:** DP-1, t_(7)_=3.7, **p < 0.01; Light phase, t_(7)_=2.7, *p < 0.05; DP-2, ns. **NREM:** DP-1, t_(7)_=4.2, **p < 0.01; Light phase, t t_(7)_=2.95,*p < 0.05; DP-2, ns. **REM:** No significant differences in any phase. Data are presented as individual paired values.

## Discussion

Elucidating how E2 regulates sleep is critical to our understanding of why women are more susceptible to sleep disruptions compared to men, particularly during hormonal transitions such as puberty, pregnancy, and menopause. This study begins to elucidate the neurobiological mechanisms underlying E2 effects on sleep-wake behavior. Our findings confirm that E2 alone reliably promotes wakefulness and suppresses NREM and REM sleep during the dark phase. Importantly, local manipulations within the MnPO demonstrate that this region is a key site of estrogenic action. Cyclo-E2 infusions were sufficient to increase wake and reduce NREM, and local infusion of the ER antagonist ICI attenuated systemic E2 effects. However, these manipulations did not fully recapitulated systemic effects of E2, particularly with respect to REM sleep. In contrast, systemic activation of both ERα and ERβ together fully reproduced the effects of E2, whereas activation of either receptor alone was insufficient. Taken together, these results highlight the MnPO as a central node where estrogen receptor signaling shapes wake and NREM regulation, while also suggesting that coordinated ER activity in additional sites may be required for the E2’s full effects on sleep-wake behavior, including REM suppression. By identifying the MnPO as a central site of E2 action, this work provides the beginnings of a neuroanatomical and molecular framework for understanding how ovarian hormones contribute to sleep disruptions across the female lifespan, offering potential targets for treating sleep disorders associated with changes in the ovarian steroid milieu.

Exogenously administered E2 has been well-documented in rodent models to suppress sleep and promote wakefulness, particularly during the dark/active phase^29–42^. This effect is also observed in female rats with endogenous ovarian hormones, where the most pronounced changes in sleep-wake behavior occur on proestrus, coinciding with peak circulating E2 levels^31,33,41,72,73^. Our findings align with these established effects of E2 on sleep regulation. However, what is not well understood is whether progesterone, which also peaks on proestrus, has additional effects on sleep-wake behavior. The few existing studies report mixed results. In female rats with prolonged exposure to E2 and progesterone via implanted capsule, E2 alone increases wake and decreases REM sleep and the combination of E2 and progesterone results in additive effects further increasing wakefulness and reducing REM sleep^36^. An early study using acute injection of E2 and progesterone also found additional effects over E2 alone^30^. These discrepancies may stem from methodological differences in dose and route of administration, as well as timing of progesterone relative to the peak of E2.

Unlike the previous studies, our exogenous E2/progesterone replacement paradigm more closely mimics the natural rise of ovarian steroid over the estrous cycle^45,46^. Our current findings support the conclusion that progesterone has no additional effect on sleep-wake behavior, reinforcing E2 as the primary driver of sleep-wake changes. However, it is important to note that progesterone administration at high doses has been shown to enhance sleep in postmenopausal women^74,75^. This effect is likely mediated by its metabolite, allopregnanolone, a known GABA-A receptor agonist, as GABA-A agonists replicate this sleep-promoting effect, while GABA-A antagonists block it.^76,77^

E2 altered sleep–wake architecture by decreasing the number of wake, NREM, and REM bouts in the dark phase and shifting bout distribution toward longer wake episodes at the expense of shorter sleep bouts. This mirrors observations in proestrus females, where wake bouts are longer and NREM/REM bouts shorter than in other cycle phases.^41^ Together with evidence from sleep deprivation studies showing ovarian steroids promote recovery sleep,^36,39,45^ these findings suggest that E2 promotes wake continuity while reducing sleep stability. Complementing these architectural effects, E2 also significantly reduced NREM-SWA, consistent with reports that SWA is lowest during proestrus, particularly at the light-to-dark transition.^31^ In contrast Swift et al. (2019) indicate that NREM-SWA may increase in specific brain regions during the dark phase of proestrus, suggesting potential regional differences in the effects of E2 on sleep homeostasis.^41^ A finding that would support multiple sites of E2 action that modulated sleep-wake behavior and physiology.

In our current study as well as our previous work, effects of E2 on sleep-wake behavior required at least 12 to 24 hours to manifest^78,79^, suggesting a genomic mechanism mediated by nuclear estrogen receptors. Given that ERα and ERβ are the primary nuclear estrogen receptors expressed in the CNS, we sought to determine their specific contributions to E2’s regulation of sleep-wake states. Selective activation of either receptor alone with PPT (ERα agonist) or DPN (ERβ agonist) failed to alter sleep-wake behavior, whereas co-administration of PPT and DPN fully replicated systemic E2, increasing wake and reducing both NREM and REM during the dark phase. These findings potentially suggest that ERα and ERβ act in concert to modulate sleep-wake behavior, rather than exerting independent or redundant effects.^80^ While prior studies have shown that individual estrogen receptors mediate other physiological functions, such as ERα’s predominant role in regulating energy balance and ERβ’s involvement in anxiety-like behaviors^51–53,81,82^, our findings highlight a potential cooperative interaction between these receptors in sleep regulation and is a focus of on-going studies.

To further investigate potential sites of E2 action, we examined ER expression in the MnPO and VLPO, two preoptic area nuclei implicated in sleep-wake control.^43^ ERα protein levels were markedly higher in the MnPO compared to the VLPO, consistent with our prior work showing that E2 influences adenosine signaling in this region.^42^ Females exhibited significantly more ERα-positive cells than males, a difference that may help explain the lack of E2-induced sleep changes in males.^40^

Although validated antibodies for ERβ were unavailable at the time of this study, fluorescent in situ hybridization confirmed the presence of both *ESR1* and *ESR2* mRNA in the MnPO. *ESR1* expression was far greater than *ESR2* in both sexes. Females again showed higher expression of both *ESR1* and *ESR2,* suggesting enhanced estrogenic sensitivity relative to males. A small subset of MnPO neurons co-expressed both receptors. While these dual-expressing cells could contribute to the cooperative ERα/ERβ requirement identified in our systemic agonist experiments, their limited numbers make it unlikely that they fully account for E2’s effects. This relatively low level of ERβ expression further supports the view that additional estrogen-sensitive sites contribute to the systemic profile of sleep–wake regulation.

This interpretation is consistent with our MnPO manipulations. Local infusion of Cyclo-E2 was sufficient to transiently increase wake and suppress NREM, and local ER antagonism with ICI attenuated but did not abolish systemic E2 effects. Importantly, neither manipulation significantly altered REM sleep, suggesting that MnPO signaling accounts for only part of the estrogenic regulation of sleep–wake behavior.

While our findings suggest a genomic mechanism, we cannot rule out the involvement of fast-acting estrogen receptors such as membrane-associated ERα, ERβ and GPR30/GPER, which may also contribute to E2 regulation of sleep-wake states, though their roles remain unclear.^83,84^.

Given the widespread distribution of estrogen receptors in the brain, identifying the precise neural circuits through which ERα and ERβ regulate sleep-wake behavior remains a critical challenge. Our previous work identified the MnPO as a potential site for estrogenic regulation of sleep-wake.^42,85^ The MnPO serves as a key integrator of homeostatic sleep signals and contains diverse neuronal populations that project to arousal-promoting regions such as the lateral hypothalamus and brainstem nuclei.^43,44,86,87^ While GABAergic in the MnPO are widely accepted as promoters of NREM sleep by inhibiting arousal pathways, recent studies show that glutamatergic MnPO neurons also contribute to sleep regulation, particularly by limiting stress-induced sleep disruption.^88^ Taken together with the current study, these findings establish the MnPO as a critical node within sleep-regulatory networks and strongly suggest that it is an important locus for E2 action.

### Limitations and Future Directions

Rodents provide a useful bioassay for studying E2 effects on sleep. Despite differences in sleep architecture between rodent and humans the neurocircuitry and sleep regulation mechanisms share key similarities, allowing for pharmacological studies. Nevertheless, in the current study as well as the other published reports on the effects of E2 on sleep-wake, it is clear that in rodents, E2 promotes the consolidated bouts of wakefulness during the dark at the expense of sleep. This may appear paradoxical to the sleep-promoting effects reported in the clinical studies of perimenopausal/menopausal women. This difference may be due to species-specific sleep patterns and/or chronotype-dependent effects of E2 (reviewed in^1^).

One limitation of the current study is that doses for pharmacological manipulations were selected from published literature rather than derived from full dose–response experiments. This raises the possibility that the doses of PPT and DPN were suboptimal when administered individually. While this possibility cannot be completely excluded, these concentrations have been effective in other behavioral paradigms, and in our study, single agonist administration was tested across both low and high doses (Supplemental Figure 5). Across the two doses no effects were detected, suggesting that receptor-specific requirements, rather than dosing alone, likely account for the absence of single agonist effects.

Another potential limitation is that local infusions into the MnPO may have spread beyond the intended target, potentially producing off-target effects. To help mitigate this, the infusion rate was slow (1µl over 10 minutes). While this possibility cannot be excluded, misplaced injection sites from the ER antagonist experiment (ICI), suggest that the off-target effects were limited as the sleep-wake behavior in these animals more closely resembled that of the controls.

Finally, the time course of estradiol’s effects differed between systemic and local delivery. Systemic administration of lipid-soluble E2 produced effects lasting ∼48 hours, whereas local infusion of water-soluble cyclodextrin-E2 yielded a shorter effect (∼18 hours). These differences likely reflect solubility, concentration, and distribution, with systemic E2 engaging broader networks and MnPO infusions restricting action to a more localized circuit.

To our knowledge, this study is the first to demonstrate that E2 signaling in the MnPO is modulates sleep-wake behavior. However, the MnPO is just one of several brain nuclei involved in the sleep-wake circuitry (for review^43,89^). It is not known whether E2 acts in other brain nuclei that are involved in sleep and/or arousal. However, estrogen receptors have been reported in various nuclei involved in sleep and arousal.^90–94^ Further research is needed to elucidate the precise cellular mechanisms through which E2 influences sleep-wake regulation, as well as its interactions with other neuromodulatory systems.

## Conclusion

Women experience significantly more sleep disruptions than men, often tied to fluctuations in ovarian hormones across puberty, pregnancy, and menopause. Despite this, the mechanisms by which E2 regulates sleep in the female brain have remained largely unexplored. This study identifies the MnPO as a key site of estrogenic action where E2 signaling is both necessary and sufficient to regulate sleep-wake behavior. Together these findings provide a beginning framework for understanding how ovarian steroid fluctuations contribute to sleep disturbances in women. Given the heightened risk of insomnia and other sleep disorders in women, these insights lay the groundwork for developing therapies to improve sleep health in women across the lifespan.

## Preprint Repositories

Some of the early preliminary data was submit to the preprint server BioRxiv

## Data Availability Statement

The data underlying this article will be shared on reasonable request to the corresponding author.

---

**Supplemental Figure 1: Estradiol had a moderate to no effect on sleep-wake behavior in adult female rats during the light phase.**

**(A)** *Right, Hourly durations for Wake, NREM and REM.* Ovariectomized (OVX) rats received systemic oil and E2 injections across three consecutive days according to our standard paradigm (Figure 2A).

**(A)** *Left*, Hourly means of wake NREM, and REM during the light phase. No consistent differences were observed across the light hours. *Right*, Boxplots of total 12-h light phase durations. On Day 3 (10 µg E2), wake decreased by approximately 20 minutes compared to oil baseline, with a similar concomitant increase in NREM. REM sleep was unchanged.

(B) Hourly means of NREM-SWA during the light phase. E2 did not alter NREM-SWA, although a robust time-of-day effect was observed.

<u>Statistical analyses:</u>

(A) *Left* (hourly means), Two-way repeated measures ANOVA (Treatment × Time, ZT).

**Wake**: Main effect of treatment F_(1.9, 30.6)_=7.6, p<0.05; Main effect of time F(_5.2, 83.9)_=76.7, p<0.0001); no interaction.

**NREM**: Main effect of treatment, F_(1.5, 25.2)_=7.75, p<0.05; Main effect of time, F_(5.5, 88.75)_=72.9, p < 0.0001); no interaction.

**REM:** no main effects or interaction.

Post hoc (Dunnett’s): On Day 3, a single hour showed decreased wake and increased NREM compared to oil baseline.

(A) *Right* (12-h light phase totals): One-way repeated measures ANOVA.

**Wake:** F_(1.9, 30.65)_=9.6, p<0.001

**NREM:** F_(1.5, 25.2)_=7.7, p<0.005

**REM:** no significant effects.

Post hoc (Dunnett’s): Wake was decreased and NREM increased following both the low (Day 2, 5 µg) and high (Day 3, 10 µg) doses of E2 compared to oil baseline. REM was unchanged. (*p<0.05), (**p<0.01)

(B) (NREM-SWA hourly means): Mixed-effects model (Treatment × ZT). Main effect of time: F_(3.7, 59.6)_=91.9, p<0.0001

All p values were adjusted for multiple comparisons. For box plots, all data points are shown; the line represents the median and the + represents the mean.

**Supplemental Figure 2: Sustained Effects of Estradiol on Sleep–Wake Behavior (Day 4).**

The rat estrous cycle is characterized by a peak in estradiol (E2) followed by a rise in progesterone on proestrus. In this model, circulating E2 levels remain elevated 24 h after the final injection, providing an opportunity to assess whether the effects of E2 persist into this window (see Figure 2A). To confirm persistence of E2 actions, sleep– wake states were assessed on Day 4 (D4) and compared with baseline (D1) and the last injection day (D3). E2-treated animals exhibited sustained increases in wake and decreases in NREM and REM during the dark phase on D4, and these values were not different from those on D3.

<u>Statistical analyses:</u>

(A) *Left* (hourly means, dark phase): Mixed-effects model (Treatment × Time, ZT).

**Wake:** Main effect of treatment, F_(1.7, 15.3)_=12.7, p<0.001; Main effect of time, F_(4.0, 35.5)_=9.6, p <0.0001; Interaction, F_(5.5, 49.7)_=3.6, p<0.01.

**NREM:** Main effect of treatment, F_(1.7, 15.6)_=10.5, p<0.005; Main effect of time, F_(3.7, 33.7)_=10.8, p<0.0001; Interaction, F_(5.7, 51.0)_= 3.4, p<0.01.

**REM:** Main effect of treatment, F_(1.5, 13.2)_= 13.3, p< 0.005; Main effect of time, F_(4.2, 38.2)_=4.5, p<0.005; Interaction, F_(5.3, 47.4)_=3.0, p < 0.05.

Post hoc (Tukey’s): On both D3 and D4, wake was increased and NREM and REM decreased at ZT14 (**p<0.005) and ZT15 (**p<0.005) compared to baseline (D1). There were no significant differences between D3 and D4.

(A), *Right* (12-h dark phase totals): Mixed-effects model.

Wake: F_(1.7, 23.0)_=12.5, p<0.0005

NREM: F_(1.7, 15.6)_=10.1, p<0.005

REM: F_(1.5, 13.2)_=13.3, p<0.005

**Post hoc (Tukey’s):** On both D3 and D4, wake was significantly increased (**p<0.01) and both NREM and REM significantly decreased (NREM *p < 0.05; REM, D3 **p<0.01 and D4 *p<0.05) compared to baseline. There were no significant differences between D3 and D4.

All p values were adjusted for multiple comparisons. For line graphs, data are presented as mean ± SEM. For box plots, all data points are shown; the line represents the median and the + represents the mean.

**Supplemental Figure 3: Progesterone in the presence of E2 had no significant effect on sleep-wake states compared to E2-alone.**

In the rat estrous cycle, a sharp rise in progesterone follows the peri-ovulatory estradiol (E2) peak. Because circulating E2 levels remain elevated 24 h after the last injection in this model, this time point (Day 4) provides an opportunity to mimic the natural proestrus progesterone surge. To test whether progesterone contributes to E2’s effects on sleep–wake behavior, E2-treated animals were divided into Post-E2 and E2 + Progesterone groups. Hourly means of wake, NREM, and REM were assessed across both the light and dark phases (grey box). Sleep– wake behavior did not differ between Post-E2 and E2 + Progesterone groups in either phase. Both groups showed sustained increases in wake and decreases in NREM and REM during the dark phase compared with baseline.

<u>Statistical analyses:</u>

Hourly means, 24h across light and dark: Mixed-effects model (Treatment × Time, ZT).

**Wake:** Main effect of treatment, F_(2, 372)_= 8.2, p<0.0005; Main effect of time, F_(5.7, 194.3)_=9.1, p <0.0001; interaction, F_(11.5, 194.3)_= 2.7, p<0.01.

**NREM:** Main effect of treatment, F_(2, 370)_=6.7, p<0.005; Main effect of time, F_(11, 370)=_10.4, p <0.0001; interaction, F_(22, 370)_=2.7, p<0.0001.

**REM:** Main effect of treatment, F_(2, 370)_=6.2, p <0.01; Main effect of time, F_(5.3, 162.1)_=4.5, p<0.001; interaction, F_(10.5, 162.1)_=2.4, p<0.05.

**Post hoc (Tukey’s):** In both Post-E2 and E2 + Progesterone groups, wake was increased at ZT13 (*p <0.05), ZT14 (***p <0.0005), and ZT15 (*p <0.05) compared to baseline (D1). Corresponding decreases were observed in NREM at ZT13 (**p <0.001), ZT14 (***p0.0005), and ZT15 (**p< 0.005). REM was decreased at ZT14 (**p <0.005) and ZT15 (*p <0.05) compared to baseline.

All p values were adjusted for multiple comparisons. For line graphs, data are presented as mean ± SEM.

**Supplemental Figure 4: Light-phase sleep architecture is minimally affected by estradiol.**

To determine whether estradiol alters sleep architecture, bout number and mean bout duration were assessed in 6 h bins during the light phase at baseline (Day 1) and following E2 treatment (Day 3; see Figure 2A). Compared to baseline, E2 reduced the number of wake and NREM bouts during the early light phase (ZT0–5), but REM bout number was unaffected. Mean bout duration for wake, NREM, and REM did not differ significantly between conditions.

<u>Statistical analyses</u>

(A) Bout number, light phase, 6h bins: Two-way repeated measures ANOVA (Treatment × Time).

**Wake:** Main effect of treatment, F_(1,16)=_9.1, p<0.01.

**NREM:** Main effect of treatment, F_(1,16)=_9.9, p<0.01; interaction, F_(1,16)=_10.6, p<0.01.

REM: No significant effects.

**Post hoc (Šídák’s):** Wake bout number was reduced (*p<0.05) and NREM bout number was reduced (***p<0.0005) during the early light phase (ZT0–5) on Day 3 compared to baseline.

(B) Mean bout duration, light phase, 6h bins: No significant effects of treatment were observed for wake, NREM, or REM.

All p values were adjusted for multiple comparisons. Data are presented in box plots with all data points. The line represents the median and the + represents the mean.

**Supplemental Figure 5. Time course of sleep–wake behavior following selective ER agonist treatments.**

OVX rats were treated with selective ERα (PPT) or ERβ (DPN) agonists, alone or in combination, alongside E2 and vehicle controls. Agonists were administered in a stepwise fashion, with a low dose on Day 1, a high dose on Day 2, and recordings continued through Day 3, the post-injection period 24 h after the final treatment (Figure 2A). Sleep–wake durations for Wake, NREM, and REM were analyzed across all three treatment days (D1–D3) in the dark phase. Only E2 and the combined PPT+DPN treatment produced significant changes between the low-dose and post-injection days, with increased wake and reduced NREM and REM. In contrast, PPT or DPN alone did not differ across days.

<u>Statistical analyses</u>

Two-way repeated measures ANOVA (Treatment × Day) with Tukey’s multiple comparisons test.

**Wake:** Main effect of treatment, F_(4,31)_=3.5, p<0.05; Main effect of day, F_(2,59.1)_=6.9, p<0.005; interaction, F_(7.6, 59.1)_=2.0, p=05.

**NREM:** Main effect of treatment, F_(4,31)_=2.6, p=0.051; Main effect of day, F_(2,59.1)_=5.5, p<0.01; interaction, F_(7.8, 60.6)_=2.0, p=0.06.

**REM:** Main effect of treatment, F(_4,31_)=6.7, p<0.0005; Main effect of day, F_(1.5,45.8)_=8.8, p<0.005; interaction, F_(6,45.8)_=3.1, p<0.05.

**Post hoc (Tukey’s):** Significant differences were detected between D1 and D3 in the E2 and PPT+DPN groups, with increased wake and reduced NREM and REM (*p<0.05, **p<0.01). No differences were observed for PPT or DPN alone.

All p values were adjusted for multiple comparisons. Data are presented in box plots with all data points; the line represents the median and the + represents the mean.

**Supplemental Figure 6: Representation of MnPO injection sites.**

**(A)** *Representative photomicrographs of cannulation sites (arrows) and stylet placement (asterisk).* **(B)** *Placement map of injection sites.* Grey circles represent correct stylet placement, and a circled x represents a miss placement. *ac, anterior commissure; 3v, third ventricle;* Scale bar =100µm.

**Supplemental Figure 7. MnPO estrogen receptor signaling does not influence sleep–wake behavior in the light phase.**

Ovariectomized (OVX) rats received oil or E2 injections combined with local MnPO infusions of vehicle (Veh) or the estrogen receptor antagonist ICI 182,780 (Fulvestrant; ICI; 50 ng/µl) in a within-animal double crossover design (see Figure 2B). Sleep–wake durations were analyzed during the light phase. No significant differences were observed in wake, NREM, or REM between Oil/Veh and Oil/ICI, nor between E2/Veh and E2/ICI. All p values were adjusted for multiple comparisons. Data are presented in box plots with all data points; the line represents the median and the + represents the mean.

**Supplemental Figure 8. MnPO estrogen receptor signaling does not regulate sleep– wake behavior in males.**

Adult male rats were implanted with transmitters and MnPO cannulas. In a within-animal design, animals first received a Veh infusion followed by an ICI infusion 25 h later, with EEG/EMG recordings collected for the 24 h following each infusion. After completing the first arm, animals were orchidectomized (ORDX), allowed to recover, and then underwent the same infusion treatments in the absence of testicular hormones. Sleep–wake durations were compared across conditions. No significant differences in wake, NREM, or REM were observed in intact or ORDX males following Veh or ICI infusions, indicating that MnPO estrogen receptor signaling does not play a major role in male sleep–wake behavior.

**Supplemental Figure 9: Representation of MnPO injection sites.**

*Placement map of injection sites.* Grey circles represent correct stylet placement, and a circled x represents a miss placement. Correct cannula placement was confirmed in 80% of cases, with misplacements excluded from the analysis.

## References

1. Mong JA, Cusmano DM. Sex differences in sleep: impact of biological sex and sex steroids. Philosophical Transactions of the Royal Society B: Biological Sciences. 2016; 371 (1688): 20150110.

2. Shechter A, Varin F, Boivin DB. Circadian variation of sleep during the follicular and luteal phases of the menstrual cycle. Sleep. 2010; 33 (5): 647–656.

3. Parry BL, Mendelson WB, Duncan WC, Sack DA, Wehr TA. Longitudinal sleep EEG, temperature, and activity measurements across the menstrual cycle in patients with premenstrual depression and in age-matched controls. Psychiatry research. 1989; 30 (3): 285–303.

4. Shechter A, Boivin DB. Sleep, Hormones, and Circadian Rhythms throughout the Menstrual Cycle in Healthy Women and Women with Premenstrual Dysphoric Disorder. International journal of endocrinology. 2010; 2010: 259345.

5. Joffe H, Massler A, Sharkey KM. Evaluation and management of sleep disturbance during the menopause transition. Semin Reprod Med. 2010; 28 (5): 404–421.

6. Dzaja A, Arber S, Hislop J, et al. Women’s sleep in health and disease. Journal of psychiatric research. 2005; 39 (1): 55–76.

7. Mendez M, Radtke RA. Interactions between sleep and epilepsy. Journal of clinical neurophysiology : official publication of the American Electroencephalographic Society. 2001; 18 (2): 106–127.

8. Pasic Z, Smajlovic D, Dostovic Z, Kojic B, Selmanovic S. Incidence and types of sleep disorders in patients with stroke. Medicinski arhiv. 2011; 65 (4): 225–227.

9. Petit D, Gagnon JF, Fantini ML, Ferini-Strambi L, Montplaisir J. Sleep and quantitative EEG in neurodegenerative disorders. J Psychosom Res. 2004; 56 (5): 487–496.

10. Redline S, Kump K, Tishler PV, Browner I, Ferrette V. Gender differences in sleep disordered breathing in a community-based sample. Am J Respir Crit Care Med. 1994; 149 (3 Pt 1): 722–726.

11. Somers VK, White DP, Amin R, et al. Sleep apnea and cardiovascular disease: an American Heart Association/american College Of Cardiology Foundation Scientific Statement from the American Heart Association Council for High Blood Pressure Research Professional Education Committee, Council on Clinical Cardiology, Stroke Council, and Council On Cardiovascular Nursing. In collaboration with the National Heart, Lung, and Blood Institute National Center on Sleep Disorders Research (National Institutes of Health). Circulation. 2008; 118 (10): 1080–1111.

12. Vock J, Achermann P, Bischof M, et al. Evolution of sleep and sleep EEG after hemispheric stroke. J Sleep Res. 2002; 11 (4): 331–338.

13. Ayas NT, White DP, Manson JE, et al. A prospective study of sleep duration and coronary heart disease in women. Arch Intern Med. 2003; 163 (2): 205–209.

14. Gangwisch JE, Heymsfield SB, Boden-Albala B, et al. Short sleep duration as a risk factor for hypertension: analyses of the first National Health and Nutrition Examination Survey. Hypertension. 2006; 47 (5): 833–839.

15. Patel SR, Ayas NT, Malhotra MR, et al. A prospective study of sleep duration and mortality risk in women. Sleep. 2004; 27 (3): 440–444.

16. Cappuccio FP, Stranges S, Kandala NB, et al. Gender-specific associations of short sleep duration with prevalent and incident hypertension: the Whitehall II Study. Hypertension. 2007; 50 (4): 693–700.

17. Meier-Ewert HK, Ridker PM, Rifai N, et al. Effect of sleep loss on C-reactive protein, an inflammatory marker of cardiovascular risk. J Am Coll Cardiol. 2004; 43 (4): 678–683.

18. Krystal AD. Depression and insomnia in women. Clin Cornerstone. 2004; 6 Suppl 1B: S19–28.

19. Lyytikainen P, Rahkonen O, Lahelma E, Lallukka T. Association of sleep duration with weight and weight gain: a prospective follow-up study. Journal of sleep research. 2011; 20 (2): 298–302.

20. Ferrie JE, Shipley MJ, Cappuccio FP, et al. A prospective study of change in sleep duration: associations with mortality in the Whitehall II cohort. Sleep. 2007; 30 (12): 1659– 1666.

21. Miller MA, Kandala NB, Kivimaki M, et al. Gender differences in the cross-sectional relationships between sleep duration and markers of inflammation: Whitehall II study. Sleep. 2009; 32 (7): 857–864.

22. Kronholm E, Laatikainen T, Peltonen M, Sippola R, Partonen T. Self-reported sleep duration, all-cause mortality, cardiovascular mortality and morbidity in Finland. Sleep medicine. 2011; 12 (3): 215–221.

23. Johnson EO, Roth T, Schultz L, Breslau N. Epidemiology of DSM-IV insomnia in adolescence: lifetime prevalence, chronicity, and an emergent gender difference. Pediatrics. 2006; 117 (2): e247–256.

24. Merklinger-Gruchala A, Ellison PT, Lipson SF, Thune I, Jasienska G. Low estradiol levels in women of reproductive age having low sleep variation. European journal of cancer prevention : the official journal of the European Cancer Prevention Organisation. 2008; 17 (5): 467–472.

25. Sharkey KM, Crawford SL, Kim S, Joffe H. Objective sleep interruption and reproductive hormone dynamics in the menstrual cycle. Sleep Med. 2014; 15 (6): 688–693.

26. Shechter A, Lesperance P, Ng Ying Kin NM, Boivin DB. Nocturnal polysomnographic sleep across the menstrual cycle in premenstrual dysphoric disorder. Sleep Med. 2012; 13 (8): 1071–1078.

27. Baker FC, O’Brien LM, Armitage R. Sex differences and menstrual-related changes in sleep and circadian rhythms. In: Kryger M, Roth T, Dement WC, eds. Principles and Practice of Sleep Medicine (5th Ed). St. Louis, Missouri: Elsevier Saunders.; 2010: 1562–1571.

28. Mong JA, Baker FC, Mahoney MM, et al. Sleep, rhythms, and the endocrine brain: influence of sex and gonadal hormones. J Neurosci. 2011; 31 (45): 16107–16116.

29. Colvin GB, Whitmoyer DI, Sawyer CH. Circadian sleep-wakefulness patterns in rats after ovariectomy and treatment with estrogen. Experimental neurology. 1969; 25 (4): 616–625.

30. Branchey M, Branchey L, Nadler RD. Effects of estrogen and progesterone on sleep patterns of female rats. Physiology & behavior. 1971; 6 (6): 743–746.

31. Schwierin B, Borbely AA, Tobler I. Sleep homeostasis in the female rat during the estrous cycle. Brain Res. 1998; 811 (1-2): 96–104.

32. Deurveilher S, Cumyn EM, Peers T, Rusak B, Semba K. Estradiol replacement enhances sleep deprivation-induced c-Fos immunoreactivity in forebrain arousal regions of ovariectomized rats. American journal of physiology Regulatory, integrative and comparative physiology. 2008; 295 (4): R1328–1340.

33. Hadjimarkou MM, Benham R, Schwarz JM, Holder MK, Mong JA. Estradiol suppresses rapid eye movement sleep and activation of sleep-active neurons in the ventrolateral preoptic area. Eur J Neurosci. 2008; 27 (7): 1780–1792.

34. Mashoodh R, Stamp JA, Wilkinson M, Rusak B, Semba K. Lack of estradiol modulation of sleep deprivation-induced c-Fos in the rat brain. Physiol Behav. 2008; 95 (4): 562–569.

35. Paul KN, Turek FW, Kryger MH. Influence of sex on sleep regulatory mechanisms. J Womens Health. 2008; 17 (7): 1201–1208.

36. Deurveilher S, Rusak B, Semba K. Estradiol and progesterone modulate spontaneous sleep patterns and recovery from sleep deprivation in ovariectomized rats. Sleep. 2009; 32 (7): 865–877.

37. Paul KN, Laposky AD, Turek FW. Reproductive hormone replacement alters sleep in mice. Neuroscience letters. 2009; 463 (3): 239–243.

38. Schwartz MD, Mong JA. Estradiol suppresses recovery of REM sleep following sleep deprivation in ovariectomized female rats. Physiology & Behavior. 2011; 104 (5): 962–971.

39. Deurveilher S, Seary ME, Semba K. Ovarian hormones promote recovery from sleep deprivation by increasing sleep intensity in middle-aged ovariectomized rats. Horm Behav. 2013; 63 (4): 566–576.

40. Cusmano DM, Hadjimarkou MM, Mong JA. Gonadal steroid modulation of sleep and wakefulness in male and female rats is sexually differentiated and neonatally organized by steroid exposure. Endocrinology. 2014; 155 (1): 204–214.

41. Swift KM, Keus K, Echeverria CG, et al. Sex differences within sleep in gonadally intact rats. Sleep. 2020; 43 (5).

42. Smith PC, Phillips DJ, Pocivavsek A, et al. Estradiol influences adenosinergic signaling and nonrapid eye movement sleep need in adult female rats. Sleep. 2022; 45 (3): 2021.2005.2027.445868.

43. Scammell TE, Arrigoni E, Lipton JO. Neural Circuitry of Wakefulness and Sleep. Neuron. 2017; 93 (4): 747–765.

44. Donlea JM, Alam MN, Szymusiak R. Neuronal substrates of sleep homeostasis; lessons from flies, rats and mice. Curr Opin Neurobiol. 2017; 44: 228–235.

45. Schwartz MD, Mong JA. Estradiol modulates recovery of REM sleep in a time-of-day-dependent manner. Am J Physiol Regul Integr Comp Physiol. 2013; 305 (3): R271– 280.

46. Holder MK, Hadjimarkou MM, Zup SL, et al. Methamphetamine facilitates female sexual behavior and enhances neuronal activation in the medial amygdala and ventromedial nucleus of the hypothalamus. Psychoneuroendocrinology. 2010; 35 (2): 197–208.

47. Holder MK, Mong JA. Methamphetamine enhances paced mating behaviors and neuroplasticity in the medial amygdala of female rats. Hormones & Behavior. 2010.

48. Holder MK, Veichweg SS, Mong JA. Methamphetamine-enhanced female sexual motivation is dependent on dopamine and progesterone signaling in the medial amygdala. Horm Behav. 2015; 67: 1–11.

49. Stauffer SR, Coletta CJ, Tedesco R, et al. Pyrazole ligands: structure-affinity/activity relationships and estrogen receptor-alpha-selective agonists. J Med Chem. 2000; 43 (26): 4934–4947.

50. Meyers MJ, Sun J, Carlson KE, Marriner GA, Katzenellenbogen BS, Katzenellenbogen JA. Estrogen receptor-beta potency-selective ligands: structure-activity relationship studies of diarylpropionitriles and their acetylene and polar analogues. J Med Chem. 2001; 44 (24): 4230–4251.

51. Harris HA, Katzenellenbogen JA, Katzenellenbogen BS. Characterization of the biological roles of the estrogen receptors, ERalpha and ERbeta, in estrogen target tissues in vivo through the use of an ERalpha-selective ligand. Endocrinology. 2002; 143 (11): 4172–4177.

52. Santollo J, Wiley MD, Eckel LA. Acute activation of ER alpha decreases food intake, meal size, and body weight in ovariectomized rats. Am J Physiol Regul Integr Comp Physiol. 2007; 293 (6): R2194–2201.

53. Patisaul HB, Burke KT, Hinkle RE, Adewale HB, Shea D. Systemic administration of diarylpropionitrile (DPN) or phytoestrogens does not affect anxiety-related behaviors in gonadally intact male rats. Horm Behav. 2009; 55 (2): 319–328.

54. Paxinos G, Watson C. The rat brain in stereotaxic coordinates. Amsterdam; Boston: Elsevier Academic Press; 2005.

55. Lu J, Bjorkum AA, Xu M, Gaus SE, Shiromani PJ, Saper CB. Selective Activation of the Extended Ventrolateral Preoptic Nucleus during Rapid Eye Movement Sleep. J Neurosci. 2002; 22 (11): 4568–4576.

56. Howell A, Osborne CK, Morris C, Wakeling AE. ICI 182,780 (Faslodex): development of a novel, “pure” antiestrogen. Cancer. 2000; 89 (4): 817–825.

57. Wakeling AE, Bowler J. ICI 182,780, a new antioestrogen with clinical potential. J Steroid Biochem Mol Biol. 1992; 43 (1-3): 173–177.

58. Wakeling AE, Dukes M, Bowler J. A potent specific pure antiestrogen with clinical potential. Cancer Res. 1991; 51 (15): 3867–3873.

59. Canal Castro C, Pagnussat AS, Orlandi L, et al. Coumestrol has neuroprotective effects before and after global cerebral ischemia in female rats. Brain Res. 2012; 1474: 82–90.

60. Lynch JF, 3rd, Winiecki P, Vanderhoof T, Riccio DC, Jasnow AM. Hippocampal cytosolic estrogen receptors regulate fear generalization in females. Neurobiol Learn Mem. 2016; 130: 83–92.

61. Rivera HM, Eckel LA. Activation of central, but not peripheral, estrogen receptors is necessary for estradiol’s anorexigenic effect in ovariectomized rats. Endocrinology. 2010; 151 (12): 5680–5688.

62. Stinson C, Logan SM, Bellinger LL, Rao M, Kinchington PR, Kramer PR. Estradiol Acts in Lateral Thalamic Region to Attenuate Varicella Zoster Virus Associated Affective Pain. Neuroscience. 2019; 414: 99–111.

63. Zurkovsky L, Brown SL, Korol DL. Estrogen modulates place learning through estrogen receptors in the hippocampus. Neurobiol Learn Mem. 2006; 86 (3): 336–343.

64. Santollo J, Torregrossa AM, Eckel LA. Estradiol acts in the medial preoptic area, arcuate nucleus, and dorsal raphe nucleus to reduce food intake in ovariectomized rats. Horm Behav. 2011; 60 (1): 86–93.

65. Santollo J, Daniels D. Anorexigenic effects of estradiol in the medial preoptic area occur through membrane-associated estrogen receptors and metabotropic glutamate receptors. Horm Behav. 2019; 107: 20–25.

66. Packard MG, Teather LA. Intra-hippocampal estradiol infusion enhances memory in ovariectomized rats. Neuroreport. 1997; 8 (14): 3009–3013.

67. Sinopoli KJ, Floresco SB, Galea LA. Systemic and local administration of estradiol into the prefrontal cortex or hippocampus differentially alters working memory. Neurobiol Learn Mem. 2006; 86 (3): 293–304.

68. Merchenthaler I, Lane MV, Numan S, Dellovade TL. Distribution of estrogen receptor alpha and beta in the mouse central nervous system: in vivo autoradiographic and immunocytochemical analyses. Journal of Comparative Neurology. 2004; 473 (2): 270–291.

69. Wilson ME, Rosewell KL, Kashon ML, Shughrue PJ, Merchenthaler I, Wise PM. Age differentially influences estrogen receptor-alpha (ERalpha) and estrogen receptor-beta (ERbeta) gene expression in specific regions of the rat brain. Mechanisms of Ageing & Development. 2002; 123 (6): 593–601.

70. Shughrue PJ, Merchenthaler I. Distribution of estrogen receptor beta immunoreactivity in the rat central nervous system. The Journal of comparative neurology. 2001; 436 (1): 64–81.

71. Morishita M, Higo S, Iwata K, Ishii H. Sex and interspecies differences in ESR2- expressing cell distributions in mouse and rat brains. Biology of sex differences. 2023; 14 (1): 89.

72. Kostin A, Alam MA, Siegel JM, McGinty D, Alam MN. Sex- and Age-dependent Differences in Sleep-wake Characteristics of Fisher-344 Rats. Neuroscience. 2020; 427: 29–42.

73. Toth A, Keseru D, Petho M, et al. Sleep and local field potential effect of the D2 receptor agonist bromocriptine during the estrus cycle and postpartum period in female rats. Pharmacol Biochem Behav. 2024; 239: 173754.

74. Schussler P, Kluge M, Yassouridis A, et al. Progesterone reduces wakefulness in sleep EEG and has no effect on cognition in healthy postmenopausal women. Psychoneuroendocrinology. 2008; 33 (8): 1124–1131.

75. Schussler P, Kluge M, Adamczyk M, et al. Sleep after intranasal progesterone vs. zolpidem and placebo in postmenopausal women - A randomized, double-blind cross over study. Psychoneuroendocrinology. 2018; 92: 81–86.

76. Lancel M, Faulhaber J, Holsboer F, Rupprecht R. Progesterone induces changes in sleep comparable to those of agonistic GABAA receptor modulators. American Journal of Physiology. 1996; 271 (4 Pt 1): E763–772.

77. Lancel M, Faulhaber J, Holsboer F, Rupprecht R. The GABA(A) receptor antagonist picrotoxin attenuates most sleep changes induced by progesterone. Psychopharmacology (Berl). 1999; 141 (2): 213–219.

78. Vasudevan N, Pfaff DW. Non-genomic actions of estrogens and their interaction with genomic actions in the brain. Frontiers in Neuroendocrinology. 2008; 29 (2): 238–257.

79. McEwen BS, Alves SE. Estrogen actions in the central nervous system. Endocrine Reviews. 1999; 20 (3): 279–307.

80. Si X, Guo T, Wang Z, et al. Neuroimaging evidence of glymphatic system dysfunction in possible REM sleep behavior disorder and Parkinson’s disease. NPJ Parkinsons Dis. 2022; 8 (1): 54.

81. Harris HA. Estrogen receptor-beta: recent lessons from in vivo studies. Mol Endocrinol. 2007; 21 (1): 1–13.

82. Miller CK, Meitzen J. No detectable changes in anxiety-related and locomotor behaviors in adult ovariectomized female rats exposed to estradiol, the ERβ agonist DPN or the ERα agonist PPT. Horm Behav. 2023; 152: 105363.

83. Arterburn JB, Prossnitz ER. G Protein-Coupled Estrogen Receptor GPER: Molecular Pharmacology and Therapeutic Applications. Annu Rev Pharmacol Toxicol. 2023; 63: 295–320.

84. Zabinska M, Wisniewska K, Wegrzyn G, Pierzynowska K. Exploring the physiological role of the G protein-coupled estrogen receptor (GPER) and its associations with human diseases. Psychoneuroendocrinology. 2024; 166: 107070.

85. Smith PC, Mong JA. Neuroendocrine Control of Sleep. Curr Top Behav Neurosci. 2019; 43: 353–378.

86. McGinty D, Gong H, Suntsova N, et al. Sleep-promoting functions of the hypothalamic median preoptic nucleus: inhibition of arousal systems. Archives italiennes de biologie. 2004; 142 (4): 501–509.

87. Alam MN, Kumar S, Bashir T, et al. GABA-mediated control of hypocretin- but not melanin-concentrating hormone-immunoreactive neurones during sleep in rats. J Physiol (Lond). 2005; 563 (2): 569–582.

88. Machado NLS, Todd WD, Kaur S, Saper CB. Median preoptic GABA and glutamate neurons exert differential control over sleep behavior. Curr Biol. 2022; 32 (9): 2011–2021 e2013.

89. Saper CB, Fuller PM. Wake-sleep circuitry: an overview. Curr Opin Neurobiol. 2017; 44: 186–192.

